# Crowd-sourced benchmarking of single-sample tumour subclonal reconstruction

**DOI:** 10.1101/2022.06.14.495937

**Authors:** Adriana Salcedo, Maxime Tarabichi, Alex Buchanan, Shadrielle M.G. Espiritu, Hongjiu Zhang, Kaiyi Zhu, Tai-Hsien Ou Yang, Ignaty Leshchiner, Dimitris Anastassiou, Yuanfang Guan, Gun Ho Jang, Kerstin Haase, Amit G. Deshwar, William Zou, Imaad Umar, Stefan Dentro, Jeff A. Wintersinger, Kami Chiotti, Jonas Demeulemeester, Clemency Jolly, Lesia Sycza, Minjeong Ko, PCAWG-11 Working Group, SMC-Het Participants, David C. Wedge, Quaid D. Morris, Kyle Ellrott, Peter Van Loo, Paul C. Boutros

## Abstract

Tumours are dynamically evolving populations of cells. Subclonal reconstruction algorithms use bulk DNA sequencing data to quantify parameters of tumour evolution, allowing assessment of how cancers initiate, progress and respond to selective pressures. A plethora of subclonal reconstruction algorithms have been created, but their relative performance across the varying biological and technical features of real-world cancer genomic data is unclear. We therefore launched the ICGC-TCGA DREAM Somatic Mutation Calling -- Tumour Heterogeneity and Evolution Challenge. This seven-year community effort used cloud-computing to benchmark 31 containerized subclonal reconstruction algorithms on 51 simulated tumours. Each algorithm was scored for accuracy on seven independent tasks, leading to 12,061 total runs. Algorithm choice influenced performance significantly more than tumour features, but purity-adjusted read-depth, copy number state and read mappability were associated with performance of most algorithms on most tasks. No single algorithm was a top performer for all seven tasks and existing ensemble strategies were surprisingly unable to outperform the best individual methods, highlighting a key research need. All containerized methods, evaluation code and datasets are available to support further assessment of the determinants of subclonal reconstruction accuracy and development of improved methods to understand tumour evolution.

## Introduction

Tumours evolve from normal cells through sequential acquisition of somatic mutations. These mutations occur probabilistically, influenced by the cell’s chromatin structure and both endogenous and exogenous mutagenic pressures^1^. If specific mutations provide a selective advantage to a cell, then its descendants can expand within their local niche. This process can repeat over years or decades until a population of cells with a common set of somatic mutations (a clone) exhibits multiple hallmarks of cancer^2,3^. Throughout this time, different tumour cell subpopulations (subclones) can emerge through drift or selective pressures across the population^4^.

Tumour evolutionary features are increasingly recognized to have clinical implications. Genetic heterogeneity has been associated with worse outcomes, larger numbers of mutations, and therapy-resistance^5–8^. The evolutionary timing of individual driver mutations influences the fraction of cancer cells that will be affected by therapies targeting them. The specific pattern of mutations and their timing can shed light on tumour aetiology and sometimes predicts therapeutic sensitivity^9^.

The process of inferring quantitative features of an individual tumour’s (sub)clonal composition based on mutational features of its genome is called subclonal reconstruction^10^, and is an common approach to quantify aspects of tumour evolution. Numerous algorithms have been developed for this task based on allelic frequencies of somatic single nucleotide variants (SNVs) and copy number aberrations (CNAs). Many apply Bayesian inference^11–14^, but a broad variety of strategies have been developed^15–17^.

Subclonal reconstruction results can vary dramatically from algorithm to algorithm^18^. Little is known about how tumour characteristics and technical parameters – such as depth of sequencing or accuracies of variant and copy-number calls – quantitatively influence the performance of subclonal reconstruction algorithms. It has even been unclear how best to quantify algorithm accuracy^19^. There is a clear need to identify which subclonal reconstruction algorithms most accurately infer specific evolutionary features, and what aspects of both the cancer itself and of the DNA sequencing experiment most influence accuracy.

To address these questions, we applied a validated framework for simulating and scoring evolutionarily realistic cancers^19^ in a crowd-sourced benchmarking challenge to quantify the accuracy of 31 strategies for subclonal reconstruction against 51 extensively annotated tumour phylogenies. Using this library of interchangeable methods, we quantify algorithm performance and show that only a small number of specific tumour features strongly influence reconstruction accuracy. These results and resources will improve the application of existing subclonal reconstruction methods and support algorithm enhancement and development.

## Results

### Challenge design

To benchmark methods for tumour subclonal reconstruction, we built upon the ICGC-TCGA DREAM Somatic Mutation Calling Challenges and their tumour simulation framework (**Figure 1a**)^19–21^. We designed 51 tumour phylogenies (**Supplementary Figure 1**) to cover wide ranges of tree topologies, tumour purities, mutation burdens and effective read depth (**Figure 1b**). Twenty-five of these phylogenies were based on manually curated tumours from the Pan-Cancer Analysis of Whole Genomes consortium (PCAWG)^22^, while 16 were based on non-PCAWG tumours^13,23–28^. The remaining ten were designed as variations of a single breast tumour, each testing a specific edge-case or assumption of subclonal reconstruction algorithms (**Extended Data Figure 1a**; ^13^). We supplemented these with a five-tumour titration series at 8x, 16x, 32x, 64x and 128x coverage^19^. For each tumour design, we simulated normal and tumour BAM files using BAMSurgeon^19^, then used GATK MuTect^29^ to identify somatic SNVs and Battenberg^13^ to identify somatic CNAs and estimate tumour purity. These were provided as inputs to participating groups, who were blinded to all other details of the tumour genome and evolutionary history.

**Figure 1.**
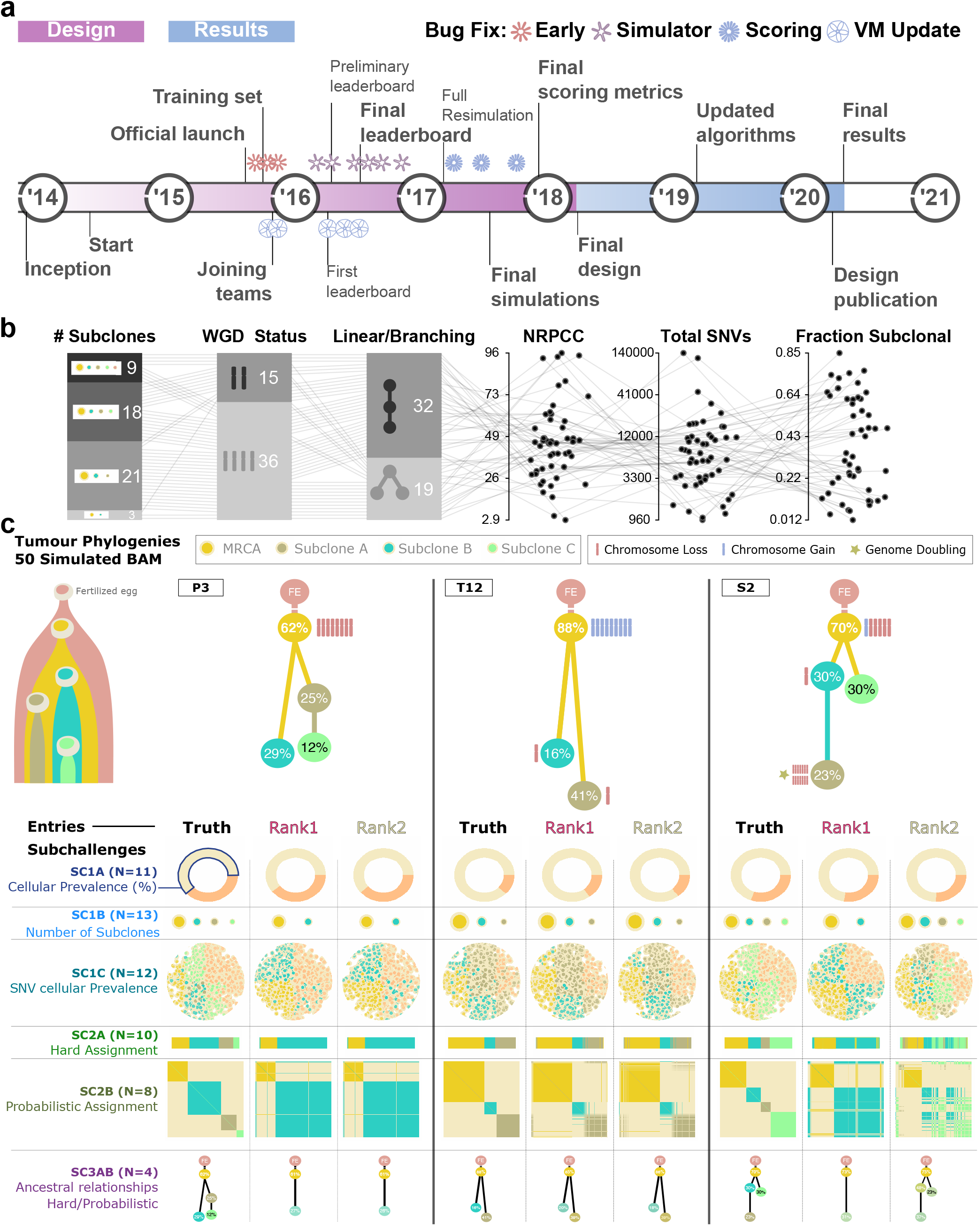
Design of the challenge **a)** Timeline of the SMC-Het DREAM Challenge. The design phase started in 2014 with final reporting in 2021. **b)** Simulation parameter distributions across the 51 tumours. From left to right: number of subclones, whole genome doubling status, linear *vs*. branching topologies, number of reads per tumour chromosomal copies (NRPCC), total number of SNVs and fraction of subclonal SNVs. **c)** Examples of tree topologies for three simulated tumours (P3, T12, and S2). For each simulated tumour, its tree topology is shown on top of the truth (column 1) and two example methods predictions (columns 2 and 3) for each SubChallenge (rows).

Participating teams submitted 31 containerized workflows that were executed in a reproducible cloud architecture^30^. Organizers added five reference algorithms: an assessment of random chance predictions, the PCAWG “informed brute-force” clustering^31^, an algorithm that placed all SNVs in a single cluster at the variant allele frequency mode and two state of the art algorithms (SOTA1: DPClust^13^ and SOTA2: PhyloWGS^11^). Each method was evaluated on seven SubChallenges (sc) evaluating different aspects of subclonal reconstruction: sc1A infers purity, sc1B subclone number, sc1C SNV cellular frequencies, sc2A hard mutation clustering, sc2B soft mutation clustering and sc3A and sc3B infer phylogeny deterministically and probabilistically, respectively (**Figure 1c**). A library of interoperable Docker containers was generated, one for each entry. These are publicly available in Synapse (https://www.synapse.org/#!Synapse:syn2813581/files/). Each prediction was scored using an established framework, with scores normalized across methods within {Tumour, SubChallenge} tuples to range from zero to one^19^. Runs that generated errors and produced no outputs, that produced malformed outputs or that did not complete within 21 days on a compute node with at least 24 CPUs and 200 GB of RAM, were deemed failures and assigned a score of NA (2,189 runs; **Supplementary Table 1**). These failures mainly occurred for two tumours with over 100,000 SNVs. To ensure our conclusions were consistent across software versions, we executed updated versions for five algorithms (**Extended Data Figure 2**; **Supplementary Table 1**). Differences were modest (r = 0.74) but varied across SubChallenges and algorithms; updates particularly influenced assessments of subclone number (sc1B; r = 0.34). In total we considered 11,432 runs across the seven SubChallenges (**Supplementary Table 1**) and refined these to 6,758 scores after eliminating failed runs, highly correlated submissions (r > 0.75) from the same team and only considering submissions made during the initial Challenge period (**Methods**; **Supplementary Tables 2-3**).

**Table 1.**
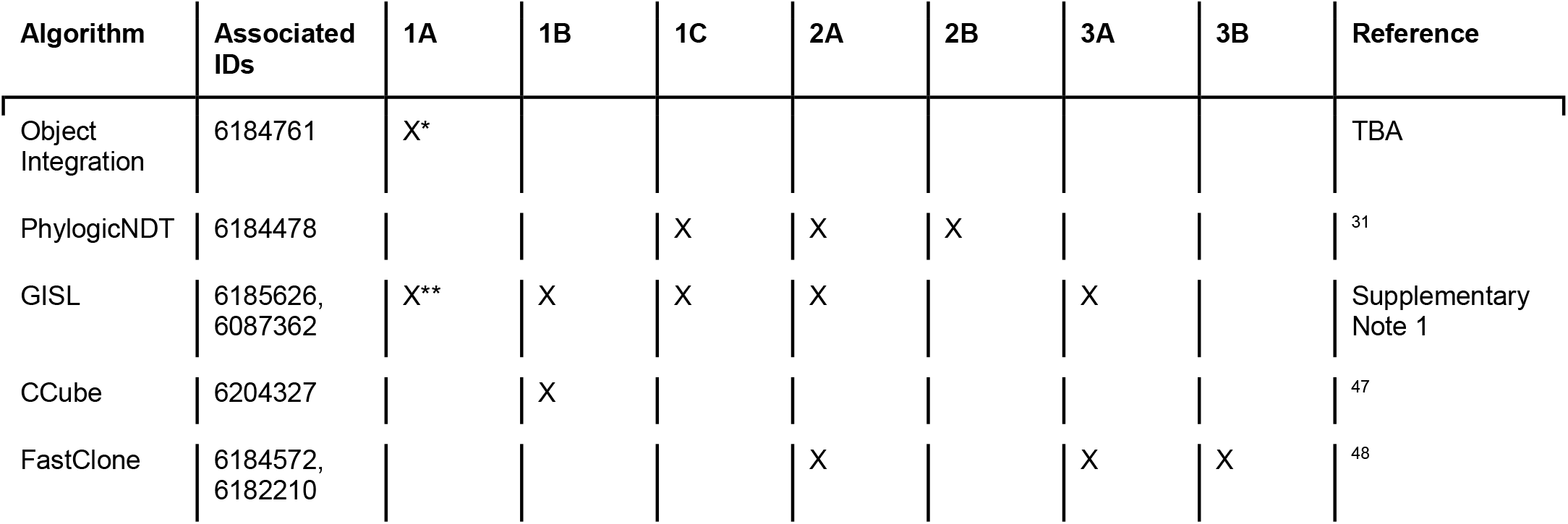
Top performing methods for each SubChallenge. SubChallenges where the method was a top performer are indicated with an X.

### Top-performing subclonal reconstruction methods

We ranked algorithms based on median scores across all tumours: no single eligible entry was the top performer across multiple SubChallenges (**Figure 2a**). For each SubChallenge, a group of algorithms showed strong and well-correlated performance (**Figure 2b, Extended Data Figure 3a-e**), suggesting multiple near-equivalent top performers. We therefore bootstrapped across tumours to test statistical significance of differences in ranks (*i*.*e*. to assess rank_entry_ < rank_best_ and assign a p-value under the null hypothesis rank_entry_ = rank_best_). sc1A and sc2B had single top-performing submissions, while two statistically indistinguishable (P > 0.1) submissions were identified for sc1B and sc1C, and three for sc2A (**Extended Data Figure 4**). The top performer for sc1A used copy number calls alone to infer purity, outperforming the purity estimates from the provided copy number calls. The second best sc1A method used a consensus of purity estimates from both copy-number and SNV clustering and was statistically indistinguishable from CNA-based purity estimates (P=0.16).

**Figure 2.**
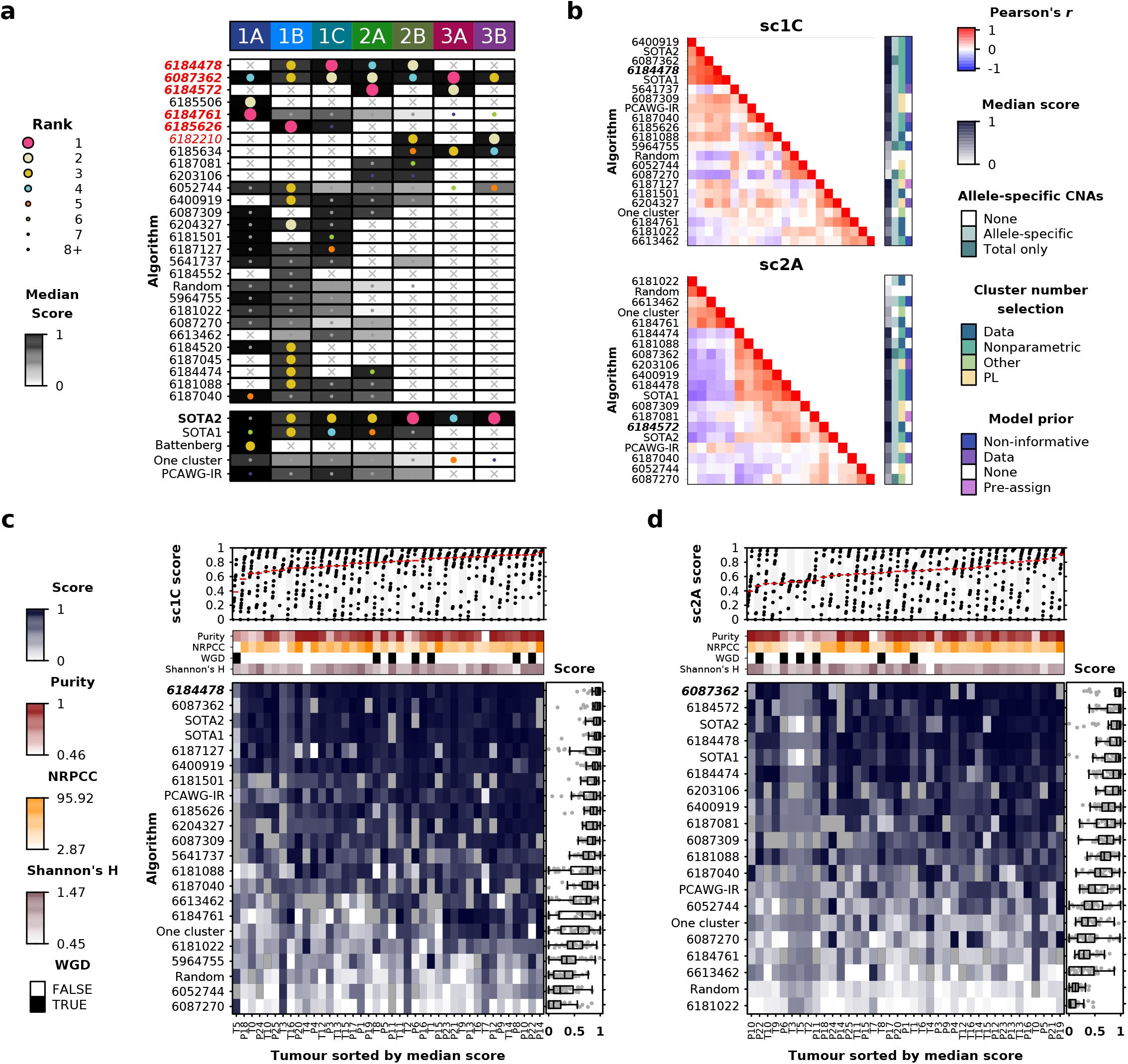
Overview of algorithm performance **a)** Ranking of algorithms on each SubChallenge based on median score. The size and colour of each dot shows algorithm rank on a given SubChallenge while the background colour reflects its median score. The top performing algorithm for each SubChallenge is shown in bold text and winning submissions are highlighted in red, italic text. **b)** Algorithm score correlations on SubChallenges 1C and 2A with select algorithm features. The top performing algorithm for each SubChallenge is shown in bold, italic text. Algorithm scores on each tumour for **c)** SubChallenge 1C and **d)** SubChallenge 2A. Bottom panels show algorithm scores for each tumour with select tumour covariates shown above. The distribution of relative ranks for each algorithm across tumours is shown in the left panel. Top panels show scores for each tumour across algorithms with the median highlighted in red. Tumours are sorted by difficulty from highest (left) to lowest (right), estimated as the median score across all algorithms.

Seven algorithms were submitted to the phylogenetic reconstruction tasks (sc3A and sc3B). Multiple algorithms were statistically indistinguishable as top performers in both challenges (**Extended Data Figure 4**), but accuracy differed widely across and within tumours. Two examples of divergent predictions are given in **Supplementary Figures 2a-b**. Predicted and true phylogenies for all tumours are at https://mtarabichi.shinyapps.io/smchet_results/, and true phylogenies are in **Supplementary Figure 1**. Algorithms differed in their ability to identify branching phylogenies (**Supplementary Figure 2c**) and in their tendency to merge or split individual nodes (**Supplementary Figure 2d**). Parent clone inference errors shared similarities across algorithms: ancestor inference for SNVs within a node was the more likely to be correct if the node was closely related to the normal (*i*.*e*. if it was the clonal node or its child) (**Supplementary Figures 2e**,**f**). When algorithms inferred the wrong parent for a given SNV, most assignment errors were to closely related nodes (**Supplementary Figure 2g**). As expected, these results emphasize that single sample phylogenetic reconstruction is most reliable for variants with higher expected alternate read counts, such as clonal variants and their direct descendants; detailed phylogenies vary widely across tumours and algorithms.

The scores of methods across SubChallenges were correlated (**Extended Data Figure 3f**). This is in part driven by patterns in the set of submissions that tackled each problem, and in part by underlying biological relationships between the problems. For example, sc1C, sc2A and sc2B assess different aspects of SNV clustering and their scores were strongly correlated with one another, but not with tumour purity estimation scores (sc1A). Rather, numerous algorithms scored highly on sc1A, suggesting different approaches are effective at estimating cellular prevalence (**Extended Data Figure 4**).

### Algorithm performance is largely invariant to tumour biology

To understand the determinants of the variability in algorithm performance between and within tumours, we considered the influence of features intrinsic to tumours. We ranked tumours by difficulty, quantified as the median score across all algorithms. We did so separately for CCF estimation (sc1C; **Figure 2c**), SNV clustering (sc2A; **Figure 2d**) and the other SubChallenges (**Extended Data Figures 3g-k**). The most and least difficult tumours differed across SubChallenges (**Supplementary Figure 3a**) and tumour ranks across SubChallenges were moderately correlated (**Supplementary Figure 3b**). SubChallenges sc2A and sc2B were the most (ρ = 0.61) while sc1C and sc3B were the least correlated (ρ = -0.10).

To determine if specific aspects of tumour biology influence reconstruction accuracy, we identified 18 plausible tumour characteristics. We supplemented these with four features that represent key experimental or technical parameters (*e*.*g*. read-depth; **Supplementary Table 2**). These 22 “data-intrinsic” features were generally poorly or moderately correlated to one another, with a few expected exceptions such as ploidy being well-correlated with whole genome duplication (WGD; **Extended Data Figure 5a)**. For each SubChallenge we assessed the univariate associations of each feature with the pool of scores from all algorithms (**Extended Data Figure 5b**). We only considered algorithms that ranked above the one-cluster solution to ensure baselines and outliers would not bias our results. As a reference we also considered Tumour ID, which captures all data-intrinsic features as a single categorical variable. We focused on the SubChallenges with large numbers of submissions and where scores can be modeled as continuous proportions using β-regression (**Methods**). Individual data-intrinsic features explained a surprisingly small fraction of the variance for sc1A, sc1C, sc2A and sc2B. Tumour ID explained ∼15% of the variance in scores, and no individual feature explained over 10%, suggesting data-intrinsic features are not exerting consistent large influences on subclonal reconstruction accuracy across algorithms.

We hypothesized that data-intrinsic features might therefore exhibit a method-specific effect that would be clearer in algorithms with generally strong performance. We repeated this univariate analysis on scores from the top five algorithms in each SubChallenge, which were moderately correlated (**Supplementary Figure 3c**). This modestly enhanced the strength of the detected associations. In sc1C the varying sensitivity of SNV detection across tumours (relative to the simulated ground truth) explained 15.7% of variance in accuracy (**Figure 3a**). In sc2A the read-depth adjusted for purity and ploidy (termed NRPCC, <u>n</u>umber of <u>r</u>eads <u>p</u>er <u>c</u>hromosome <u>c</u>opy^10^) explained 19.8% of the variance across tumours. The total number of SNVs and the number of subclonal SNVs explained 9.3% and 9.2% of the variance for sc1C, as might be expected since both define the resolution for subclonal reconstruction^10^. These results indicate that data-intrinsic features either explain little of the variability in subclonal reconstruction accuracy or do so in ways that differ widely across algorithms.

**Figure 3.**
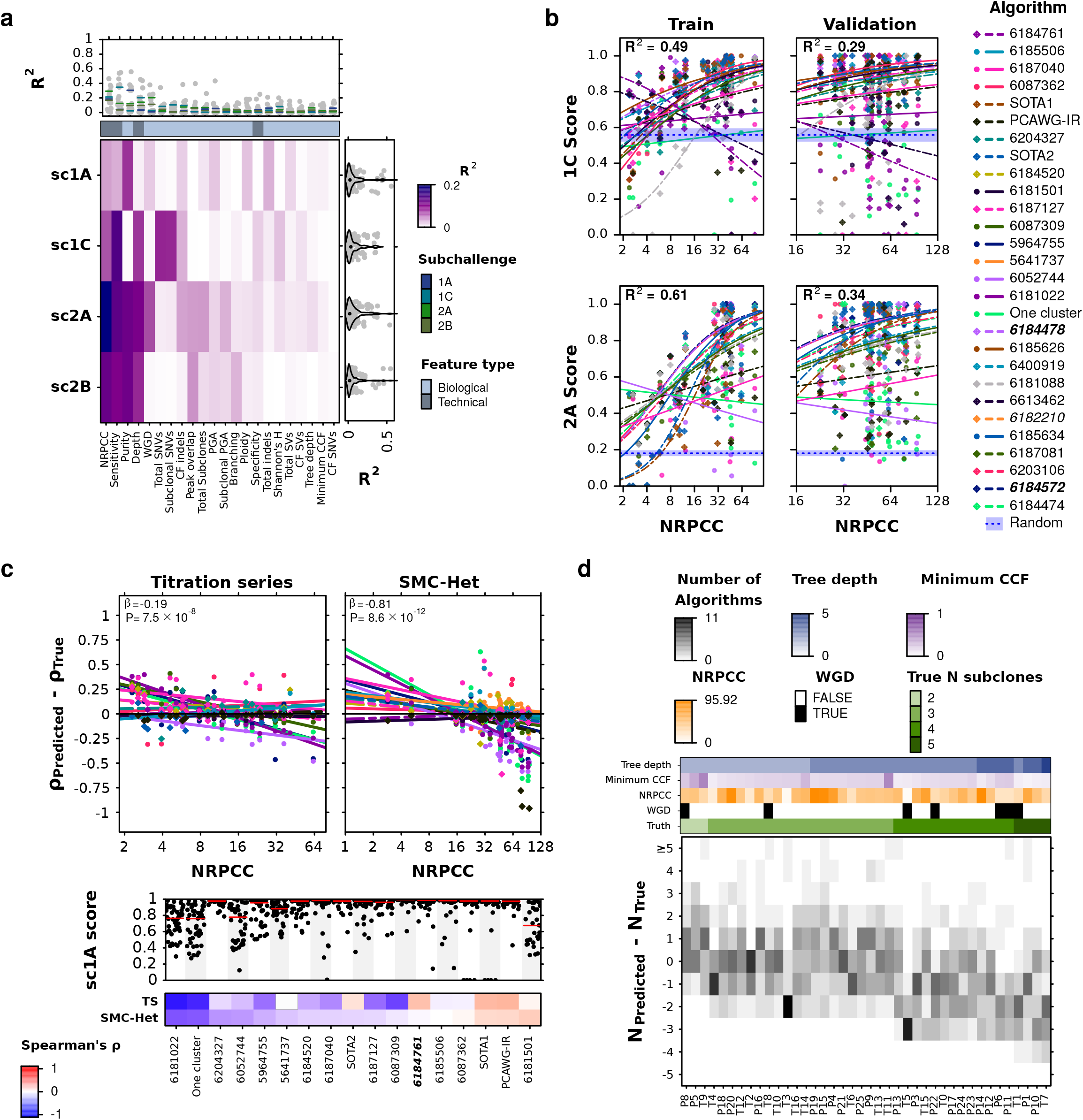
Tumour features influence subclonal reconstruction performance and biases **a)** Score variance explained by univariate regressions for the top five algorithms in each SubChallenge. Heatmap shows R^2^ for univariate regressions for features (x-axis) on SubChallenge score (y-axis) when considering only the top five algorithms. The right and upper panels show the marginal R^2^ distributions generated when running the univariate models separately on each algorithm, grouped by SubChallenge (right) and feature (upper). Lines show the median R^2^ for each feature across the marginal models for each SubChallenge. **b)** Models for NRPCC on sc1C and sc2A scores when controlling for algorithm ID. The left column shows model fit in the training set composed of titration series tumours (sampled at five depths each) and five additional tumours (N=10 individual tumours). The right column shows fit in the test set (N=30 tumours; comprising the remaining SMC-Het tumours after removing the edge cases). Blue dotted lines with a shaded region show the mean and 95% CI based on scoring ten random algorithm outputs on the corresponding tumour set. The top performing algorithm for each SubChallenge is shown in bold italic text. **c)** Effect of NRPCC on purity error. Top panels show purity error with NRPCC accounting for algorithm ID with fitted regression lines. sc1A score across tumours for each algorithm are shown in the panel below. Bottom heatmap shows Spearman’s ρ between purity error and NRPCC for each algorithm. The winning entry is shown in bold. **d)** Error in subclone number estimation by tumour. Bottom panel shows subclone number estimation error (y-axis) for each tumour (x-axis) with the number of algorithms that output a given error for a given tumour. Tumour features are shown above. NRPCC is number of reads per chromosome copy; CCF is cancer cell fractions; CF is clonal fraction (proportion of mutations in the clonal node); PGA is percent of the genome with a copy number aberration. See **Methods** for detailed descriptions of each of these.

### Algorithmic and experimental choices drive reconstruction accuracy

Given the relatively modest impact of data-intrinsic features on performance, we next focused on algorithm-intrinsic features. We first modeled performance as a function of algorithm ID, which captures all algorithmic features together. Algorithm choice alone explained 19-35% of the variance in scores in each SubChallenge (**Extended Data Figure 5c**). This exceeded the ∼15% explained by Tumour ID, despite our assessment of more tumours than algorithms.

To better understand the effect of algorithm choice, we quantified 12 specific characteristics of each algorithm. For example, we annotated whether or not each method adjusted allele frequencies for local copy-number (**Extended Data Figure 5d**). The variance explained by the most informative algorithm feature was 1.5-3x higher than that of the most informative tumour feature (**Extended Data Figure 5c**). Our analysis highlighted Gaussian noise models as particularly disadvantageous for SNV co-clustering (sc2A) relative to Binomial or Beta noise models (GLM B_gaussian_ = -0.98, P = 1.43 × 10^-15^, R^2^ = 0.11). This trend became stronger when we compared algorithms with Gaussian noise models to those with Binomial noise models and adjusted for tumour ID (B_gaussian_ = -1.11, P < 2 × 10^-16^, R^2^ = 0.35).

The strong impact of algorithm choice on performance led us to hypothesize that data-intrinsic features show algorithm-specific influences on performance. We therefore developed multivariate models to control for algorithm ID when modeling data-intrinsic features. After making this change, SNV caller sensitivity, tumour purity, and experimental read-depth were significantly associated with increased scores for nearly all SubChallenges (q < 0.05). These associations were consistent whether we analyzed all algorithms that exceeded the baseline (**Extended Data Figure 5e**) or only the top five algorithms for each SubChallenge (**Supplementary Figure 3d**). Taken together, our results show that algorithm choice is the strongest driver of subclonal reconstruction accuracy, followed by technical data-intrinsic features. Biological data-intrinsic features are weak determinants of subclonal reconstruction accuracy.

### Optimizing experimental design for subclonal reconstruction

Most data-intrinsic features reflect aspects of tumour biology not known *a priori*. In contrast, the main controllable technical feature is sequencing coverage. We investigated the sensitivity of subclonal reconstruction to this experimental design choice by considering NRPCC. By adjusting sequencing coverage for tumour purity and ploidy, NRPCC provides an excellent estimate of power in subclonal reconstruction^10^. We modeled the relationship between NRPCC and SNV co-clustering SubChallenge scores (sc1C and sc2A) using a generalized linear model in which we controlled for algorithm ID because of the strong influence of this feature in our univariate analyses above. We fit the model on five tumours with a coverage titration series (five points per tumour^19^) and on five randomly selected tumours, leading to 373 scores from these 10 tumours. We then assessed model generalizability on 466 scores from 30 tumours. Nine edge cases and two tumours with a high mutation burden (>50k SNVs) were excluded from both the training and testing cohorts. As expected, higher NRPCC increased sc1C and sc2A scores for most algorithms (**Figure 3b**). Increasing NRPCC improves co-clustering by reducing read-sampling noise, thereby improving subclone resolution^10,31^. We observed an unexpected saturation effect: at high NRPCC, most variability in scores was due to differences among algorithms. These data quantify a clear benefit to tumour sequencing to an NRPCC of at least 32 for subclonal reconstruction from a single sample across the range of algorithms tested here.

We replicated these analyses for estimation of tumour purity (sc1A). Lower NRPCC was associated with over-estimation of tumour purity (sc1A) in both the titration series and in the SMC-Het cohort (**Figure 3c**). This likely occurs because in low-coverage sequencing data, SNVs detected on a few reads are indistinguishable from background. These false negatives lead to a truncated binomial distribution and over-estimation of the average frequencies of detected SNV clusters^10,31^. Conversely, high NRPCC increases the number of subclonal mutations detected, causing some algorithms to underestimate purity (especially the naive one-cluster and random algorithms). In a similar way, NRPCC influenced the prediction of subclone number (sc1B). More algorithms underpredicted the number of subclones as the tree depth and the true subclone number increased (**Figure 3d**; B_tree depth_ = -1.18, P = 1.60 × 10^-41^, ordinal regression, likelihood ratio test), suggesting there is a limit to how many subclones can be distinguished at a given NRPCC. Consistently, the number of subclones predicted increases with NRPCC for a given tumour for most algorithms (**Extended Data Figure 6a**; B = 0.71, P = 2.99 × 10^-24^). These data emphasize that it is critical to report NRPCC and interpret estimates of tumour subclonal diversity in that context.

Finally, we asked if other tumour features might bias prediction of purity and subclone number. We used multivariate penalized regression with leave-one-out cross-validation to model sc1A and sc1B errors. After controlling for algorithm ID, the sc1A model explained 40.1% of variance and the sc1B models explained 57.1%. The multivariate model for purity estimation error highlighted that algorithms are less likely to underpredict purity as the clonal fraction (CF) of SNVs and the percent genome altered (PGA) increase, but are more likely to overestimate purity when the true purity is low (**Extended Data Figure 6b**). The subclone number error model showed algorithms are more likely to underestimate the number of subclones if there is a whole genome duplication (WGD). These results suggest that increasing power *i*.*e*. NRPCC for SNV clustering and allelic imbalance for copy-number calling, is especially important if there is *a priori* knowledge that a given tumour or tumour type is prone to low purity, clonal fraction, PGA, or is likely to harbour a WGD^10,31^. These features should be considered when interpreting subclonal reconstruction results and confirm NRPCC as a crucial study design parameter.

### Sources of error in SNV cellular prevalence estimation

Estimating what fraction of cancer cells each SNV occurs in is one of the most fundamental goals of subclonal reconstruction, shedding light on the evolution of mutational processes in a tumour^3,31–33^. To understand errors in these estimates, we focused on the 20 algorithms that produced submissions for both sc1C and sc2A. For each tumour, we annotated the SNV subclone assignments (sc2A output) with the predicted cellular prevalence for that subclone (sc1C output; **Figure 4a**). Most algorithms accurately determined whether an SNV is clonal: 14/20 had both median specificity and sensitivity above 80% (**Figure 4b**). Clonal assignment specificity increased with NRPCC as more subclonal SNVs are correctly assigned, leading to improved accuracy (**Figure 4c**; **Supplementary Figure 3a**; B_log2(nrpcc)_ = 0.29, q-value = 3.11 × 10^-17^) and decreased with SNV caller precision (B_log2(precision)_ = -1.24 q-value = 1.94 × 10^-14^, **Supplementary Figure 4a)**. Accuracy also slightly decreased with mutational burden and tumour clonal fraction (**Supplementary Figure 4a**).

**Figure 4.**
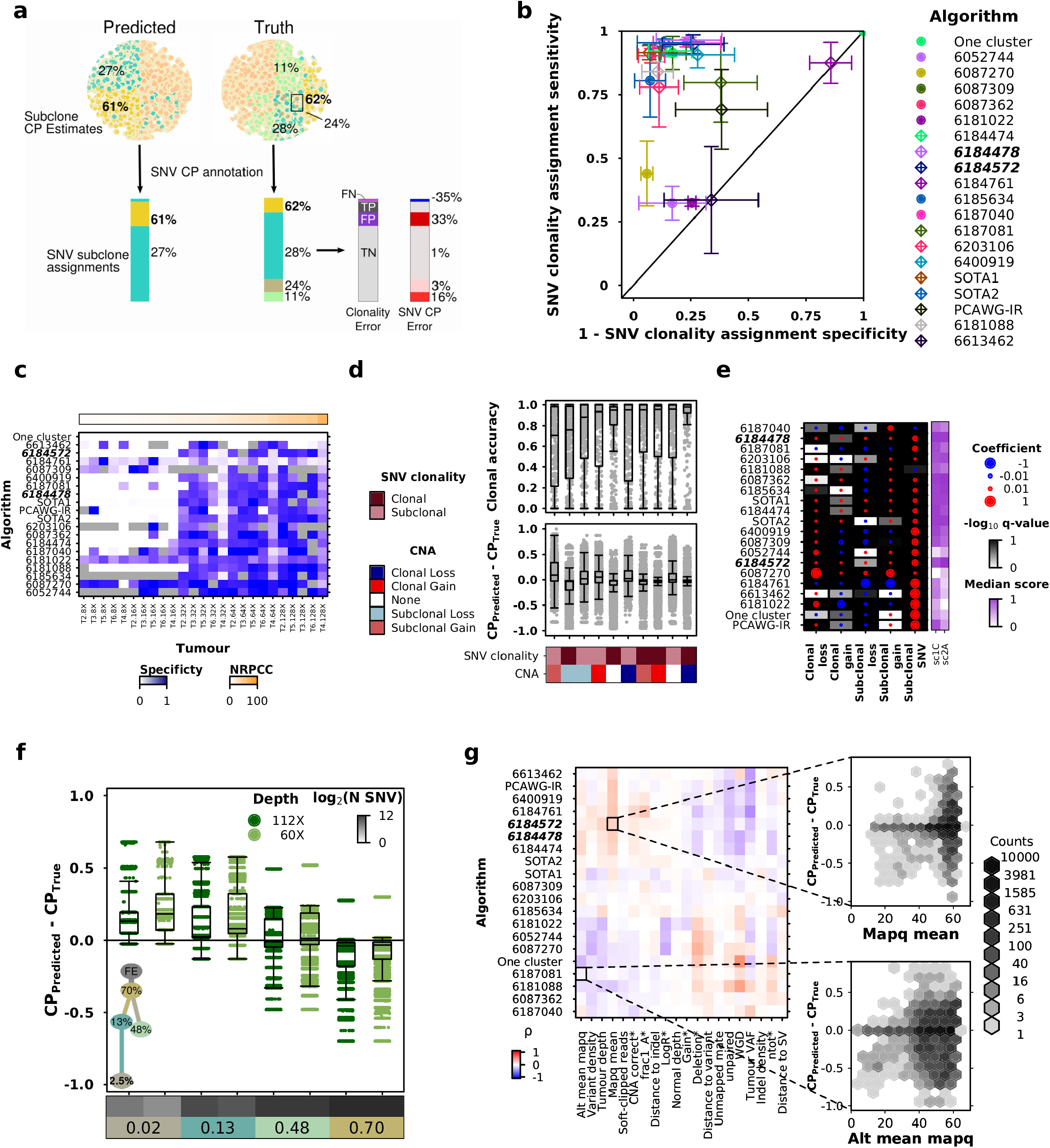
Impacts of genomic features on SNV subclonality predictions **a)** Schematic showing how outputs from SubChallenge 1C and 2A were used to annotate SNV CP for each entry. **b)** Mean clonal SNV detection sensitivity and specificity for each entry with standard errors. Winning entries for sc1C and sc2A are highlighted in bold. **c)** Clonal SNV detection F-scores for each entry on each tumour. **d)** Clonal accuracy for each entry and tumour tuple (top) and SNV CP estimation error for each entry (bottom). **e)** Effect size and FDR-adjusted p-values for entry-specific linear models for SNV CP error by CNA type and SNV clonality with median sc1C and sc2A scores. **f)** SNV CP error grouped by subclone for a corner case tumour simulated at two depths. **g)** Correlation between BAM features and Battenberg output features with SNV CP error for each entry. Only features that had an absolute correlation > 0.1 are shown. Battenberg features are noted with a star and top performing algorithms are highlighted in bold italic.

The inference of SNV clonality was impacted by the underlying copy number state of the genomic region. Subclonal CNAs significantly reduced SNV clonality assignment accuracy relative to clonal CNAs after controlling for algorithm and tumour ID (B_subclonal CNA_ = -0.21, P = 1.14 × 10^-6^, GLM). SNVs that arose clonally in a region that then experienced a subclonal loss had the least accurate clonal estimates (**Figure 4d**; B_clonal SNV x subclonal loss_ = -0.33, P = 3.06 × 10^-2^; **Supplementary Table 3**). Subclonal losses on the mutation-bearing DNA copy reduce VAF, causing many algorithms to underestimate the cellular prevalence of these SNVs (W_SNV clonal_ = 1.04 × 10^10^, P < 2.2 × 10^-16^; Wilcoxon rank-sum test for SNVs in subclonal deletions, **Supplementary Table 3**). Similarly, algorithms overestimated SNV cellular prevalence in regions with subclonal gains and subclonal SNVs (W_SNV clonal_ = 2.96 × 10^9^, P < 2.2x 10^-16^; Wilcoxon rank-sum test; **Supplementary Table 4**). This resulted in lower accuracy (B_subclonal SNV * subclonal gain_ =-0.32, P = 8.0 × 10^-3^, GLM; **Figure 4d**; **Supplementary Table 4**). Biases in CP estimation due to CNAs differed among algorithms (**Figure 4e**). To assess if robustness to CNAs impacts performance, we associated the proportion of variance in SNV CP error explained by CNA status and SNV clonality in these models with algorithm score. Algorithms whose CP estimates were more robust to CNAs better estimated the overall subclonal CP distribution (sc1C; ρ_CNA_= -0.43) and better co-clustered SNVs (sc2A; ρ_cna_= -0.37; **Supplementary Figure 4b**).

Because subclonal CNAs can be difficult to detect, we investigated whether copy number calling errors aggravated the effects of CNAs on estimation of cellular prevalence. Across all tumours, clonal CNA regions were nearly perfectly detected by our CNA caller (Battenberg), in part because they had been carefully curated during simulation (**Extended Data Figure 7a**). By contrast, 7/68 subclonal losses and 25/48 subclonal gains were entirely missed. Additionally, Battenberg inferred gains rather than losses for four CNAs and mis-estimated copy number for two subclonal gains. The titration series tumours showed that accuracy of subclonal CNA detection is strongly influenced by tumour NRPCC (**Extended Data Figure 7b**). Within a given tumour, elastic-net logistic regression showed that CNAs in low CP subclones and in SNP-poor regions were less accurately detected (**Extended Data Figure 7c**). While Battenberg CNA calling errors did not significantly impact the accuracy of SNV clonality assignment, algorithms were more likely to overestimate CP for SNVs on segments with incorrect CNA states, with consistent direction of error biases (**Extended Data Figure 7d**; **Supplementary Table 5**).

SNV features also shaped error profiles independent of CNAs. Almost all algorithms were more likely to overestimate the CP of subclonal SNVs (**Figure 4d-e**) due to reduced power at lower tumour read depths^10,13,31^. Examining two edge-case tumours with identical architectures emphasized that this bias increases for lower subclone CPs and NRPCC (**Figure 4f**). To quantify how other sources of error in SNV and CNA calls propagate to subclonal reconstruction, we derived 53 measures of variant call quality from the BAM files, VCF files and Battenberg outputs (see Methods) that we hypothesized could impact CP estimation and correlated them with CP error. Variant call quality was associated with CP error in patterns that varied across metrics and algorithms, with mean SNV mapping quality showing positive associations for many algorithms (**Figure 4g**).

### Pragmatic optimization of algorithm selection

Driven by these insights into individual algorithm performance for the seven subclonal reconstruction SubChallenges and its underlying determinants, we next sought to optimize algorithm selection across an arbitrary set of SubChallenges. To visualize algorithm performance across all SubChallenges, we projected both algorithms and SubChallenges onto the first two principal components of the scoring space, explaining 66% of total variance (**Figure 5a**). This visualization simultaneously shows algorithm performance across SubChallenges (coordinate of each algorithm on each SubChallenge axis), dissimilarities across SubChallenges (angle between SubChallenge axes) and across algorithms (distance between algorithms). The blue “decision axis” shows the axis of average score across SubChallenges when all SubChallenges are weighted equally and is stable to small fluctuations in these weights (shown by the decision “brane” around the decision axis; **Figure 5a**).

**Figure 5.**
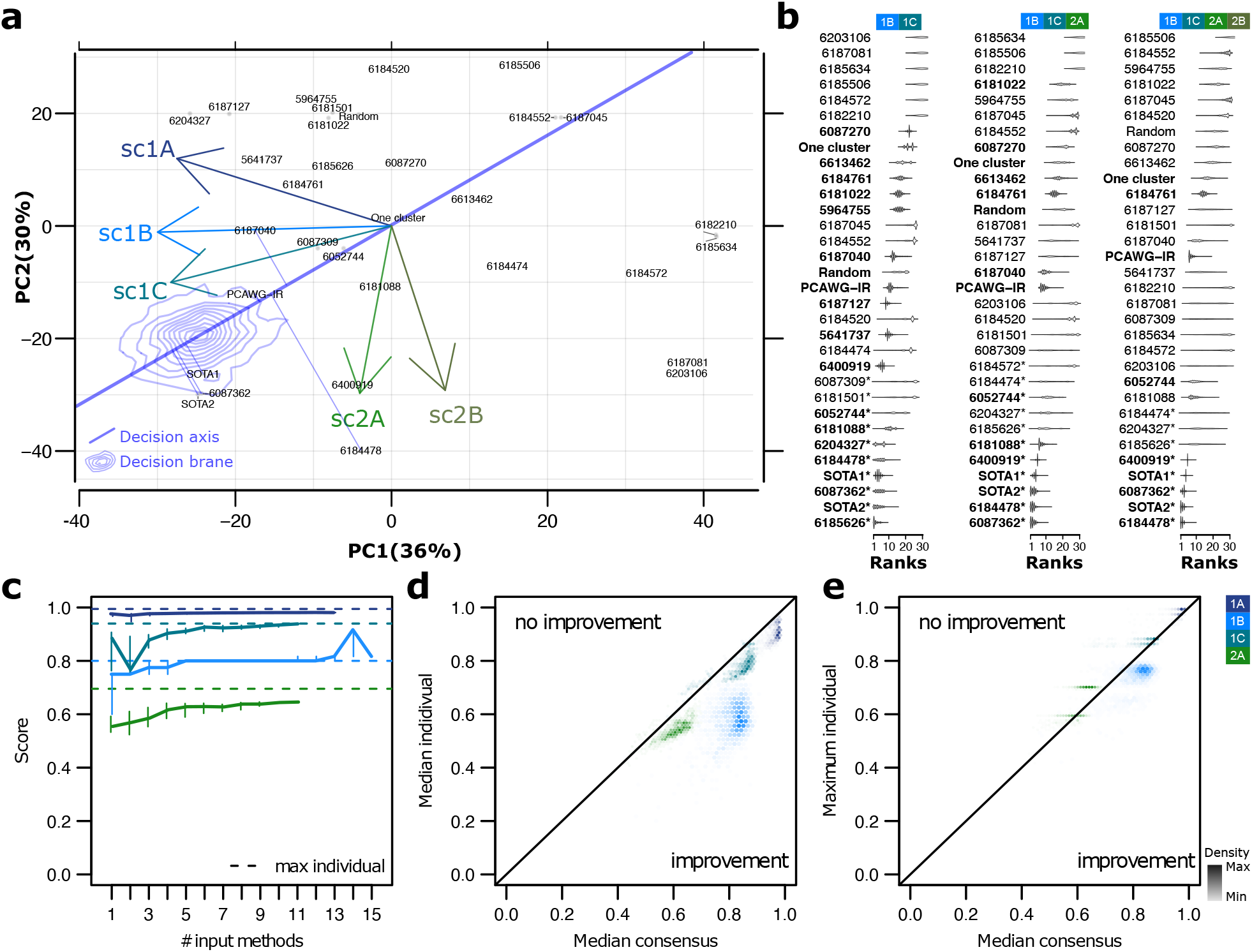
Performance across multiple algorithms and SubChallenges **a)** Projections of the algorithms and SubChallenge axes in the principal components of the score space. A decision axis is also projected and corresponds to the axis of best scores across all SubChallenges and tumours, when these are given equal weights. The five best methods according to this axis are projected onto it. A decision area in blue shows the density of decision-axes coordinates after adding random fluctuations to the weights. **b)** Rank distribution of each method from 40,000 sets of independent random uniform weights given to each tumour and SubChallenge in the overall score. From left to right: SubChallenges 1B+1C; 1B+1C+2A; 1B+1C+2A+2B. Methods in bold generate outputs for all considered SubChallenges; names of the algorithms have a star if they are ranked first at least once. **c)** Four SubChallenges for each of which one ensemble approach could be used (sc1A - median, sc1B - floor of the median, sc1C - WeMe, sc2A - CICC; **Online Methods**), we show the median and first and second tertiles (segments) of the median scores across tumours of independent ensembles based on different combinations of N methods (N varying on the x-axis). The dashed line represents the best individual score. **d)** Colour-coded hexbin densities of median ensemble vs. median individual scores across all combinations of input methods. The identity line delimiting is shown to delimit the area of improvement. **e)** Same as **d)** for maximum individual scores instead of median.

The decision axis is subjective, as it depends on arbitrary weights given to each tumour and SubChallenge. To understand the sensitivity of algorithm selection to these weights, we simulated forty thousand “studies”. In each such study, weights were randomly assigned to each SubChallenge and each tumour (**Online Methods**). We used the weighted average scores across SubChallenges and tumours to rank algorithms within each study. We performed the simulation experiments for three groups of SubChallenges: {sc1B, sc1C}, {sc1B, sc1C, sc2A}, and {sc1B, sc1C, sc2A, sc2B} (**Figure 5b**). For each algorithm, we then visualized the distribution of their ranks across all studies. Across groups of SubChallenges, 12 algorithms (35%) reach a top rank within at least one study, while 22 (65%) were never ranked first. Algorithms consistently among the top performers across SubChallenges included 6185626, 6204327, 6087362, 6184478, and the two state-of-the-art methods. Because the choice of weights is ultimately user-dependent, we created a dynamic web application for modeling the influence of different selections: https://mtarabichi.shinyapps.io/smchet_results/.

Ensemble approaches have previously been used in many different areas of biological data science to combine outputs from multiple algorithms and improve robustness^21,34,35,31^. They have not been widely explored for subclonal reconstruction, in part because many subclonal reconstruction outputs are complex and heterogeneous^31^. To begin assessing the potential of ensemble approaches to improve subclonal reconstruction, we identified and ran ensemble methods for individual SubChallenges based on median or voting approaches, which serve as conservative baselines. Specifically, we used: the median of sc1A and the floor of the median for sc1B, and the ensemble methods recently developed by the PCAWG consortium^31^ for sc1C and sc2A (**Online Methods**). We ran these ensemble methods for each SubChallenge on estimates from a large representative subset of all possible algorithm combinations (**Online Methods**) and for all tumours after excluding the ten special cases and two tumours with over 100, 000 SNVs for which only five algorithms produced outputs.

The median ensemble performance increased with the number of input algorithms for all SubChallenges (**Figure 5c**). The ensemble performance was also more consistent across tumours for sc1A and sc1B when more input algorithms were used, as shown by the decreasing variance in the scores (**Supplementary Figure 5**). Ensemble approaches outperformed the best individual methods for sc1B, but not for sc1A, sc1C and sc2A (**Figure 5c**). Rather, for these three SubChallenges, ensemble approaches had above-median performance (**Figure 5d**,**e**). In line with this, the highest ensemble performance was reached with a low number of input algorithms (two for sc1A, four for sc1B, three for sc1C and three for sc2A). These results suggest that ensemble approaches, particularly those built from multiple top-performing methods, represent a robust algorithm choice. In situations where the top-scoring algorithms for a specific dataset are not known, Dentro *et al*.^*31*^ suggested using an ensemble of all algorithms, which would also result in robust solutions. The results from our dataset nevertheless show that the proposed ensemble methods do generally improve performance over current methods despite significant computational costs. Future work on bespoke ensemble approaches for subclonal reconstruction is therefore indicated.

## Discussion

Cancer is an evolutionary process, and subclonal reconstruction from tumour DNA sequencing has become one central way this process is quantified^3,31,36,37^. Subclonal reconstruction is a complex and multi-faceted mathematical and biological process, with multiple distinct components^19^. Despite rapid proliferation of new methodologies, there has been limited benchmarking, or even surveys of the relative performance of multiple methods on a single dataset^3,10,18^. Further, despite the clear value of multi-sample and single-cell sequencing strategies, clinical studies have almost exclusively eschewed these for pragmatic, cost-effective bulk short-read sequencing of index or metastatic lesions^38,39^.

We report a crowd-sourced, unbiased benchmarking of subclonal reconstruction algorithms for single-sample designs. We show that characteristics of the experimental design (sequencing depth) and cancer types (mutation load, purity, copy number, *etc*.) influence accuracy, especially through their influence on NRPCC^10^. These results highlight important trends in the influence of the underlying copy number states on CP estimation. Algorithms are limited in the number of subclones they can confidently detect at a given depth, but their resolution increases with NRPCC. Practitioners should consider optimizing NRPCC rather than read-depth for single-sample subclonal reconstruction, and multi-region sequencing may particularly improve subclone detection sensitivity^10,40^ in low purity cancer types. Other tumour and algorithm features influence the scores in an algorithm-dependent fashion, and the choice of algorithm is the major determinant of high-quality subclonal reconstruction.

Unlike for other applications in data science such as mutation calling, current ensemble approaches for subclonal reconstruction are only as good as the best algorithms, except for identifying the number of subclones. These ensemble approaches are based on voting and averaging strategies, which might be too simplistic for sc1C and sc2A and could explain why they do not improve performance. Further development of ensemble strategies might be required to best combine the outputs of multiple algorithms and increase the performance and robustness. For these future developments, and since currently different algorithms are best at different subtasks of subclonal reconstruction, we provided online tools to extend this benchmarking, as well as a web application to help users choose the best algorithm for their dataset and question of interest (https://mtarabichi.shinyapps.io/smchet_results/).

A key opportunity for simulator improvement will be to better account for and model different aspects of cancer evolution, such as the ongoing branching evolution in the terminal (leaf) subclones^16^, the power effect and errors in SNV and CNA calling. The influence of improved simulations will also likely interact closely with specific point mutation detection strategies, suggesting future work focusing jointly on these two key algorithmic features. Recent single-cell WGS development might help build benchmarking datasets complementary to simulation-based, providing pseudo-bulk as ground truth^41–43^. Constructing an accurate ground truth from these datasets will bring its own algorithmic challenges from taking the single-cell noise into account. On-going benchmarks based on realistic and robust datasets will support continuous algorithm development, and ultimately clinical translation.

## Acknowledgments

The authors thank the members of their labs for support, and Sage Bionetworks and the DREAM Challenge organization for their ongoing support of the SMC-Het Challenge. In particular, they thank T. Norman, J.C. Bare, S. Friend and G. Stolovitzky for their technical support and scientific insight. The authors thank Google Inc. (in particular N. Deflaux) for their support of the ICGC-TCGA DREAM Somatic Mutation Calling Challenge. P.C.B. was supported by Prostate Cancer Canada and is proudly funded by the Movember Foundation - Grant #RS2014-01, by a Terry Fox Research Institute New Investigator Award, by a CIHR New Investigator Award and by the NIH through awards P30CA016042, U01CA214194, U24CA248265 and R01CA244729. Q.D.M. is supported by a Canada CIFAR AI chair through the Vector Institute and through NIH award P30CA008748. This project was supported by Genome Canada through a Large-Scale Applied Project contract to P.C.B., S.P. Shah and R.D. Morin. This work was supported by the Discovery Frontiers: Advancing Big Data Science in Genomics Research program, which is jointly funded by the Natural Sciences and Engineering Research Council (NSERC) of Canada, the Canadian Institutes of Health Research (CIHR), Genome Canada and the Canada Foundation for Innovation (CFI). This work was supported by the Francis Crick Institute, which receives its core funding from Cancer Research UK (FC001202), the UK Medical Research Council (FC001202), and the Wellcome Trust (FC001202). For the purpose of Open Access, the authors have applied a CC BY public copyright license to any Author Accepted Manuscript version arising from this submission. This project was enabled through access to the MRC eMedLab Medical Bioinformatics infrastructure, supported by the Medical Research Council (grant number MR/L016311/1). AS was supported by a CIHR Canadian Graduate Scholarship and Michael Smith Foreign Study Scholarship. MT was supported as a postdoctoral researcher of the F.R.S.- FNRS and a postdoctoral fellow by the European Union’s Horizon 2020 research and innovation program (Marie Skłodowska-Curie Grant agreement no. 747852-SIOMICS). JD was supported as a postdoctoral fellow of the European Union’s Horizon 2020 research programme (Marie Skłodowska-Curie Grant Agreement No. 703594-DECODE) and the Research Foundation– Flanders (FWO 12J6916N). PVL is a Winton Group Leader in recognition of the Winton Charitable Foundation’s support towards the establishment of The Francis Crick Institute. D.C.W. is supported by the Li Ka Shing foundation.

## Author Contributions

**Initiated Study:** PCB, QDM, PVL, DCW, KE

**Developed Methodology:** AS, MT, SE, IU, WZ, LS, MK, JD, SD, KH, CJ, AGD, JAW, HJ, KZ, TOY, DA, YG, GHJ, IL

**Data Analysis:** AS, MT, AB, KC

**Supervised Research:** PCB, PVL, QDM, KE, DCW, DA, YG, IL

**Wrote First Draft of Paper:** AS, MT, PVL, PCB

**Approved Paper:** All authors

## Figure Legends

Extended Data Figure 1. Design and scoring of special case tumours

**a)** Designs of special case tumours (top row) and their scores across SubChallenges. Each point in the strip plots represents an entry score and the red line shows the median. **b)** Heatmap of scores for sc1C and sc2A for each entry on the corner case tumours. Tumour T5 is considered as the baseline.

Extended Data Figure 2. Effects of algorithm version updates

Updated (y-axis) and original (x-axis) for five algorithms on the SMC-Het tumours. Point colour reflects the difference in the algorithm’s relative rank (r. rank) for that tumour.

Extended Data Figure 3. Overview of SubChallenge scores

**a-e)** Correlation in scores among algorithms. Each row and column is an entry for a specific SubChallenge, with colour reflecting Spearman’s ρ between entries across the main 40 SMC-Het tumours (excluding the corner cases and two tumours with > 100k SNVs where only five algorithms generated outputs), or the subset both algorithms successfully executed upon. Algorithms are clustered by correlation. Columns are sorted left-to-right in the same order that rows are top-to-bottom, thus values along the principal diagonal are all one. **f)** Correlation in scores among SubChallenges **g-k)** Scores for each tumour for SubChallenge 1A including Battenberg purity estimates as a reference **g)** sc1B **h)** sc2B **i)** sc3A **j)** and sc3B **k)** on the SMC-Het tumours. The top performing algorithm for each SubChallenge is shown in bold text and the winning submission is shown in italic. Bottom panels show algorithm scores for each tumour with select tumour covariates shown above. The distribution of relative ranks for each algorithm across tumours is shown in the left panel. Top panels show scores for each tumour across algorithms with the median highlighted in red.

Extended Data Figure 4. Rank generalizability assessment

To evaluate generalizability of ranks and differences amongst algorithms, bootstrap 95% confidence intervals were generated for median scores (left column) and ranks (right column) based on 1000 resamples. The top ranking algorithms are marked with a star for each SubChallenge and highlighted in bold on the x-axis. Winning submissions are highlighted in red. For any entry with confidence intervals overlapping those of the top ranking algorithm, bootstrap P-values comparing the rank of that algorithm to the top ranking algorithm are shown: P(rank_entry_ ≤ rank_best_). P-values for equivalent top performers (P>0.1) are highlighted in red. Algorithms are sorted by the median of their relative rank (rank/maximum rank) on each SubChallenges and top performing algorithms are highlighted in bold. Battenberg is included as a reference for sc1A.

Extended Data Figure 5. Tumour feature score associations

**a)** Correlations among tumour features and their distributions (boxplot, top). NRPCC is number of reads per chromosome copy; CCF is cancer cell fraction; CF is clonal fraction (proportion of mutations in the clonal node); PGA is percent of the genome with a copy number aberration after correcting for ploidy. See **Methods** for detailed descriptions of each. **b**,**c)** Score variance explained by univariate generalized linear models (β-regressions with a logit link) for scores generated with tumour **(b)** and algorithm **(c)** features. Models were fit on scores from all algorithms ranking above the one cluster solution on a given SubChallenge. Heatmap shows R^2^ for univariate GLMs for features (x-axis) on SubChallenge score (y-axis) on the full dataset, gray indicates missing values where models could not be run. The right and upper panels show the marginal R^2^ distributions generated when running the univariate models separately on each algorithm and tumour (for tumour and algorithm features, respectively). Tumour and algorithm ID were not included in the marginal models as the number of levels would be equivalent to the number of observations in the data subset. Lines show the median R^2^ for each feature across the marginal models for each SubChallenge. **d)** Distribution of algorithm features. **e)** Results of generalized linear models for tumour features on scores (β regression with a logit link) that controlled for algorithm-ID. The size of the dots shows the effect size and the background colour shows the P-value after FDR adjustment. Effect size interpretation is similar to that of a logistic regression, representing a one unit change in the log ratio of the score relative to its distance from a perfect score (*i*.*e*. βx=log(score/(1-score)). The bottom panel shows the results of modes fit on the full dataset. The top panel shows the same bi-variate models were fit on scores from the top five algorithms.

Extended Data Figure 6. Mutational feature error associations

**a)** Error in subclone number estimation for each algorithm on each tumour (center). Top panel plot shows NRPCC for each tumour. Right panel shows subclone number estimation error correlations with NRPCC. The top performing algorithm for SubChallenge sc1B is shown in bold italic text. **b)** Coefficient from penalized regression models for tumour features on purity estimation error (x-axis) and subclone number estimation error (y-axis).

Extended Data Figure 7. Battenberg CNA assessment

**a)** Battenberg errors for clonal and subclonal CNAs. The proportion of CNAs with correctly or incorrectly inferred clonality and copy number is shown in the heatmap. The total number of each type of CNA is indicated by the barplot on the right. **b)** Battenberg accuracy in the titration series tumours. **c)** Clonal accuracy for each entry and tumour combination (top) and SNV CP estimation error (bottom) for each entry. **d)** Effect sizes from a L1-regularized logistic regression for genomic features on Battenberg accuracy.

Supplementary Figure 1. True tumour designs

All 51 phylogenies originally designed for the challenge (52 trees are shown but T5 and S1 (shaded) are the same phylogeny based on PD4120 - this topology is both from the literature and a special case). From the fertilized egg (FE) to the first clone and subclones, we show cellular prevalences as percentages in the circles, next to which copy number events (CN) with losses (-) and gains (+) of whole chromosomes and whole-genome duplication (WGD) events are shown, along with total number of SNVs and SVs. The length of the branches are proportional to the number of SNVs.

Supplementary Figure 2. Phylogeny inference assessment

**a)** Sample true (left) and predicted tree phylogenies for T12. Each node is annotated with its CP. Branch length is proportional to the number of SNVs in a given node and the area of each colour inside corresponds to the proportion of SNVs it contains from the corresponding true node. **b)** As **a)** for the special case tumour S4 which had two clonal nodes. **c)** Each algorithm’s error profile for detecting branching phylogenies (*e*.*g*. nonlinear phylogenetic trees). **d)** The probability that each SNV from the same clone in the true tree was predicted to be in a different clone (left) and the probability that two SNVs predicted to be in the same clone are actually from different ones in the true tree. Each point represents the probability from one tumour for a given algorithm. **e)** The probability that the predicted parent of a randomly drawn SNV is correct across all tumours, with tumour specific covariates shown in the top-most heatmap. **f)** The mean probability and standard error that the predicted parent for an SNV randomly drawn from a predicted tree is correct depending on its inferred relatedness to the clonal node (*i*.*e*. 0 indicates the SNV is within the predicted clonal node and 3 indicates the SNV is within a subclone that is the great-grandchild of the predicted clonal node). **g)** For an SNV randomly drawn from a predicted tree and a SNV randomly drawn from its predicted parent clone, the mean probability and standard error of each error case across tumours for each algorithm. Algorithms are ordered by median sc3A score and the top performing algorithm for SubChallenge sc3A is shown in bold italic text.

Supplementary Figure 3. Profiling tumour difficulty and feature associations with score

**a)** Tumour rank among SubChallenges, based on median score in a given SubChallenge. Bottom panel shows relative rank (rank/total number of tumours) for each tumour on each SubChallenge and the corresponding tumour features are shown above. Relative rank for each tumour on each SubChallenge is also shown in the top scatterplot. Tumours are ordered by rank product across SubChallenges. **b)** Correlation for median tumour score among SubChallenges. **c)** Correlations in scores among the top five algorithms in each SubChallenge. The highest absolute correlation for each SubChallenge is shown. **d)** Results of univariate generalized linear models for tumour features on scores (β regression with a logit link) for each of the top five algorithms for sc1C and sc2A (sorted by ascending rank). The top-performing algorithms are highlighted in bold-italic. The size of the dots shows the feature effect size and the background colour shows the P-value for feature coefficients after FDR adjustment. Effect size interpretation is similar to that of a logistic regression, representing a one unit change in the log ratio of the score relative to its distance from a perfect score (*i*.*e*. βx=log(score/(1-score)).

Supplementary Figure 4. SNV cellular prevalence error profiling

**a)** Results of generalized linear models for tumour features on clonal accuracy (β-regression with a logit link) that controlled for entry-ID. The size of the dots shows the effect size and the background colour shows the P-value after Bonferroni adjustment. **b)** Partial residual plots showing the relationship between sc1C and sc2A scores after adjusting for covariates and the proportion of variance in SNV CP error explained by CNAs and SNV clonality for each algorithm specific model in **Figure 4e**. Lines show the mean effect averaged across all covariates and the shaded region shows the confidence interval.

Supplementary Figure 5. Variance of ensemble performance as function of number of input algorithms

For sc1A, sc1B, sc1C and sc2A, we show the variance of the ensemble method minus the lowest variance of the individual methods (y-axis) as a function of the number of input methods (on the x-axis). The dashed line represents the lowest variance of the individual methods.

## Online Methods

### Tumour designs and simulations

We designed 51 realistic tumour tree topologies with underlying subclonal structure: 16 tumour trees were inspired by published phylogenies^13,23–28^, 25 were based on manually-reconstructed PCAWG trees^22^ and 10 cases were special theoretical cases based on the highly curated PD4120^13^. Tumours from the literature and from the PCAWG study covered some of the most common cancer types (breast, prostate, lung, colorectal, and leukemia) as well as other sometimes less represented cancer types (pancreatic, sarcoma, kidney, brain, lymphoma, head and neck, thyroid), respectively (**Supplementary Table 1**).

PCAWG manual tree building was performed using DPClust (*v2*.*1*.*0*) and Battenberg (*v2*.*2*.*10*)^13^, using the pigeon-hole principle and mutation-to-mutation phasing to constrain the possible tree topologies. When multiple tree topologies were possible, we picked one at random for the simulation, while balancing branching and linear topologies across the full set of simulated tumours.

For each node, we associated a cellular prevalence, specific whole-chromosome copy-number events, a number of SNVs and SVs, as well as expected trinucleotide contexts, which all were taken as input by our simulator^19^.

As described previously^19^, we used a custom BAMSurgeon^19,21^ pipeline (implemented in Perl *v5*.*26*.*3*) to simulate BAM files with underlying tree topology and subclonal structure for the 51 tumours. Briefly, we began by aligning a high-depth (300x), Illumina paired-end publicly available BAM file (Genome in A Bottle GM24385) that was part of a father, mother, son trio using bwa (*v*.*0*.*7*.*10*) and the *hs37d5* human reference. Following a standard variant calling pipeline, we phased reads using PhaseTools (*v1*.*0*.*0*)^19^, achieving median phased contig length of ∼85kb. We then partitioned each phase and chromosome sub-BAM to simulate subclonal structure, adjusting the depth of each read pool by its cellular prevalence and total fractional copies (*i*.*e*. to simulate chromosome-length CNAs). We then spiked in SNVs, SVs, and indels into each read-pool using BAMSurgeon while preserving phylogenetic ordering (so except for deletion events, a child subclone would contain its parent’s mutations). SNVs were distributed semi-randomly to follow pre-specified trinucleotide signatures and replication timing biases. We then merged sub-BAMs across phase and chromosome to obtain the final tumour BAMs. To obtain realistic SNV calls and copy-number profiles, MuTect (*v1*.*1*.*5)*^*29*^ and Battenberg (*v2*.*2*.*10*)^13^ were run on the simulated tumour-normal BAM files.

Battenberg was run to identify clonal and subclonal copy number changes. Battenberg segments the mirrored B-allele frequencies (BAF) of phased heterozygous SNPs identified in the normal germline sample. It then selects a combination of purity and ploidy that best aligns the data to integer copy-number values in the tumour, akin to ASCAT^44^. Finally, it infers mixtures of up to two allele-specific copy number states from the BAF and logR of the obtained segments^13^. We compared the purity and ploidy values to the expected values from the designs and refitted the profiles if they did not agree. For this, we constrain a copy number state of a clonally aberrated chromosome to its known design state. Reversing ASCAT equations, we can infer ploidy and purity from a given chromosome’s BAF and logR and derive the profile using the new pair of ploidy and purity values. Only in special cases breaking the assumptions, especially those harbouring a subclonal whole-genome doubling such as PD4120, estimated purity values are not expected to closely match the design. Algorithms were run and scored on tumour VCFs and Battenberg outputs that excluded the X and Y chromosomes.

### Scoring metrics

For each SubChallenge, we used different metrics that respected a set of criteria, as described in ^19^. Here we summarize these metrics:

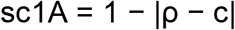

where ρ is the true cellularity, c is the predicted cellularity and |x| is the absolute value of x. Note that we require that 0 ≤ ρ ≤ 1 and 0 ≤ c ≤ 1.

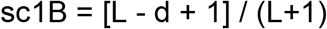

where L ≥ 1 is the true number of subclonal lineages, d is the absolute difference between the predicted and actual number of lineages, d = min(|κ - L|, L+1). We do not allow d to be higher than L+1 so that the SC1B score is always ≥ 0.

sc1C=1-EMD

where EMD is the normalized earth mover’s distance

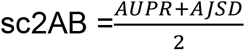

where AUPR is the normalized area under the precision recall curve and AJSD is the normalized average Jensen-Shannon divergence. We normalize AUPR and AJSD by the worst AUPR and AJSD obtained by two extreme methods, namely assigning all SNVs to one cluster and assigning each SNV to its own cluster. sc2A takes the hard assignments whereas sc2B the soft-assignment matrix.

sc3AB = PCC

where PCC is the Pearson correlation coefficient between the predicted and true values from the co-clustering matrix, cousin matrix, ancestor descendant matrix and the transposed ancestor descendant matrix. sc3A takes the hard assignments whereas sc3B the soft-assignment matrix.

### Scoring and ranking

We scored outputs obtained from participant submitted Dockerized Galaxy workflows using a Python (*v2*.*7*.*18)* implementation of the scores described above (https://github.com/uclahs-cds/tool-SMCHet-scoring). Algorithm outputs were scored against truth files based on perfect SNV calls which contained all SNVs spiked in each tumour. False negatives were added to 1c,2a,2b,3a and 3b outputs as a single cluster with a cellular prevalence of zero that was derived from the normal. False positives were excluded from outputs before scoring. We normalized scores *s* within each tumour and SubChallenge across methods using min-max normalization, *i*.*e*. offsetting and scaling such that the lowest and highest scores were set to 0 and 1, respectively:

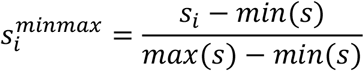

Where 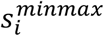 and *s*_*i*_ are the min-max normalized score and raw scores of method *i*, respectively. We normalized the titration series tumours simultaneously across all depths for a given tumour.

We ranked algorithms by normalized score across the 51 SMC-Het tumours, assigning any tied algorithms equal ranks. The best methods were defined as those with the highest median score across all tumours for which they produced a valid output.

As missing data could have been caused by technical restrictions that may not apply to users (*eg*. users would typically downsample SNVs in SNV dense tumours) and the correct penalty for missing data is subjective, we did not penalize missing outputs. However, interested users can assign scores of zero to missing outputs in the interactive app and explore how they impact algorithm rankings (https://mtarabichi.shinyapps.io/smchet_results/).

### Random methods

For sc1A, we draw a single number from a uniform distribution between 0.2 and 0.99. For sc1B, we draw from 4 integer values {1,2,3,4} with probabilities {0.2, 0.3, 0.3,0.2}, respectively. For sc1C, we assign one cluster to a CCF 1, and if there are more than one cluster, we assign random CCF values to the other clusters by drawing from a uniform distribution between 0.2 and 0.9. We then assign a random number of SNVs to each CCF cluster by drawing uniformly from 1 to 10. For sc2A, we assign a proportion of SNV per cluster by drawing uniformly from 1 to 10 for each cluster. We then randomly assign classes to SNVs. For sc2B, we generate 100 random vectors of SNV assignment to subclones and run the function *comp*.*psm* from the R package mcclust (*v1*.*0*) to obtain the proportions of co-clustering.

### Linear models for tumour and algorithm features

All statistical analyses were performed in R (*v3*.*5*). For each SubChallenge, we first removed algorithms from the same team with scores that were highly correlated across tumours (r > .75), retaining the algorithm with the highest median score for each SubChallenge. We derived 22 features to describe each tumour. Key features were defined as follows

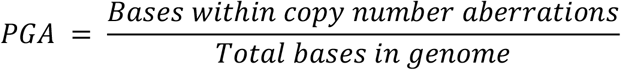

Where CNAs were defined as segments within the Battenberg output where total clonal or subclonal copy number deviated from the integer tumour ploidy.

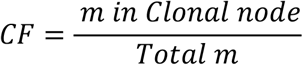

Where m is the count of SNV, Indels or SVs.

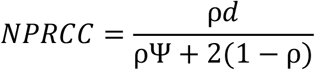

Where d is read depth, ρ is purity and Ψ is tumour ploidy.

Peak overlap was calculated by fitting density curves to each subclone in CCF space after adjusting each tumour’s variant allele-frequencies using true CNAs and cellular prevalences. To compute the relative proportion of CCF space covered by multiple subclones (peak overlap), we calculated the area underneath multiple CCF density curves relative to the total area as approximating integrals using the trapezoidal rule for each tumour. SNV, indel, and SV counts were derived from the ground-truth files used to generate each tumour.

We collected algorithm features from teams through an online form filled at the time of algorithm submission into the challenge. For each algorithm feature within each subchallenge, we removed levels represented by fewer than three algorithms as well as any labeled ‘other’ to enhance model integrity and interpretability.

We then assessed the impact of tumour and algorithm features on scores using β-regressions with the R package betareg (*v*.*3*.*2*) with a logit link function for the mean and an identity link function for Ψ (which models variance) with only an intercept term ^45^. We analyzed only sc1A, sc1C, sc2A and sc2B with β-regressions as scores for sc1B are discrete proportions (difference between the true and predicted subclone number relative to true subclone number) and measures of variance explained from binomial GLMs would not be directly comparable. Effect size interpretation is similar to that of a logistic regression, representing a one unit change in the log ratio of the expected score relative to its distance from a perfect score (*i*.*e*. βx=log(score/(1-score)). Because they represent change to a log ratio, the predicted change on a linear scale will depend on the reference score. See **Figure 3B**, for an example of effect size visualizations on a linear scale. We ran univariate models with only tumour features when we considered only the top five scoring algorithms in each SubChallenge (**Figure 3A**), as well as models that included both tumour and algorithm features when we considered all algorithms that ranked above the one cluster solution in a given SubChallenge (**Extended Data Figure 5C**). We used the same procedures to assess feature associations when controlling to algorithm ID. For these analyses we excluded corner case tumours and two tumours with >100k SNVs (P2 and P7) where only five algorithms produced outputs.

### Linear models for error bias

Bias in purity was assessed by taking the difference between the predicted and true purity for each tumour. We modeled inverse normal transformed errors using a linear regression that allowed interactions between NRPCC and algorithm ID in both the titration series and the SMC-Het tumours (excluding corner cases). As the SMC-Het tumours contained two lower NRPCC tumours, we verified results remained consistent in their absence. We then extended this analysis to multivariate modeling with elasticnet regressions as implemented in glmnet (*v*.*2*.*0-18*). Models were trained and assessed using nested cross validation where one tumour was held out in each fold. We tuned lambda and alpha in the inner loop and retained the value that achieved the lowest root mean squared error across the held out samples. In each fold, we also removed features that were >70% correlated. We used the same framework on the full dataset to train the final model. We computed R^2^ based on predictions in the held out samples of the outer loop to estimate predictive performance.

We similarly analyzed the difference between the predicted and true number of subclones. For statistical modeling we only included observations where error < 8 to minimize the effect of outliers and used a cumulative link ordinal regression implemented in MASS (*v7*.*3-51*.*6*) to model the effect of NRPCC on subclone .number estimation error when controlling for algorithm ID. We extended these to multivariate models using L1 regularized ordinal regression as implemented in ordinalNet *(v2*.*9)*. We trained and assessed these models using leave one tumour out cross validation. One tumour was held out in each fold and R^2^ was computed from correlating model predictions to the held out tumours. Within each fold we removed strongly correlated features (r >0.7) and ƛ was tuned using AIC. We report effect sizes from the final model which was trained on the full dataset. We repeated both the purity estimation error and subclone number estimation error multivariate analysis with and without algorithm ID terms. Effect sizes were congruent for both models but R^2^ decreased without algorithm ID terms.

### Genomic feature models

True CNA status was called based on the known truth. If a region experienced both clonal and subclonal CNAs, then CNAs were labeled subclonal. Genomic features were extracted from the MuTect (*v1*.*1*.*5)* VCF files using the Variant Annotation R package and from BAM files using Rsamtools (*v1*.*34*.*1*) and bam-readcount (commit 625eea2). We modeled clonal accuracy using β-regressions as described above. SNV CP error was modeled using linear regressions following an inverse normal transform. We excluded the corner case tumours from all modeling unless stated otherwise.

### Battenberg assessment

For assessing Battenberg accuracy, Battenberg copy number calls were obtained from the first solution provided in the Battenberg outputs. If a region was represented by multiple segments, we weighed each segment by its relative length and averaged their copy number estimates. We considered a clonal CNA to be correct if the total copy number for the segment matched the total true copy number of the region. Similarly, a subclonal copy number event was correct if Battenberg provided a clonal and subclonal copy number solution (P < 0.05) and the total copy number matched the true copy number of any of the tumour leaf clones (*e*.*g*. clones that did not have children). We trained and assessed the L1-regularized logistic regression for correct Battenberg CNA calls using nested cross validation as described above, tuning lambda using the inner loop. As the dataset was highly unbalanced, within each fold we sampled 250 CNAs where Battenberg was correct, and included all 104 CNAs where Battenberg was incorrect and resampled with replacement an additional 50 incorrect CNAs. Within each fold we removed correlated features (r>0.7) and optimized ƛ for sensitivity in the held out samples. We repeated this procedure on the full dataset to train the final model.

### Ensemble subclonal reconstruction

We ran ensemble methods on the outputs of four SubChallenges: sc1A, sc1B, sc1C and sc2A. For sc1A, the ensemble approach is the median of the outputs. For sc1B, it is the floor of the median. For sc1C, we ran WeMe^31^, which takes a weighted median of the CCF and the proportion of SNVs assigned to the CCF to construct a consensus location profile, and ignores individual SNVs assignments. Finally, consensus for sc2A was performed using CICC^31^, which takes the hard cluster assignment of each SNV to clusters and performs a hierarchical clustering on the co-assignment distances across methods between mutations to identify SNVs that most often cluster together across methods. We ran these approaches on 39 tumours, excluding the special cases and the two tumours with the largest number of SNVs, P2 and P7, for which most algorithms did not provide any outputs. For an increasing number of input algorithms, we ran the ensemble approaches on all possible combinations of algorithms, except when the possible number of combinations was >200, in which case we randomly sampled 200 combinations without replacement.

### Scores across multiple SubChallenges and multi-criteria decision

Akin to the PROMETHEE methodology used in decision engineering for the subjective choice of alternatives based on a set of quantitative criteria^46^, we perform principal component analyses on the weighted means of the scores across tumours in the SubChallenge dimensions, representing ∼66% of the variance in the data. We project methods and SubChallenges in that space. A decision axis is also projected which is a weighted mean of the scores across SubChallenges. Projection of the methods onto that axis leads to a method ranking. To assess the stability of the decision axis upon weight changes, we also show a density area for the decision axis projection defined by 3,000 decision axes obtained after adding from -50% to 50% changes drawn uniformly to the SubChallenge weights. We also randomly assigned weights to tumours (200 times) and SubChallenges (200 times) from uniform distributions and derived 40,000 independent rankings.

### Best performing methods

#### Data Visualization

Figures were generated using R (*v4*.*0*.*5*), Boutros Lab Plotting General (*v6*.*0*.*0*)^49^, lattice (*v0*.*20-41*), latticeExtra (*v0*.*6-28*), gridExtra (*v2*.*3*) and Inkscape (*v1*.*0*.*2*). Partial residual plots were generated with the effects package (*v4*.*2*). colour palettes were generated using the RColorBrewer package (*v1*.*1-2*).

### Accession Codes

BAM files are available in EGA at EGAS00001002092. SNV, SV, CNA, and Indel calls and corresponding truth files are available at https://www.synapse.org/#!Synapse:syn2813581/files/. Participant-submitted Docker containers are available in Synapse at https://www.synapse.org/#!Synapse:syn2813581/docker/, and Galaxy workflows at https://github.com/smc-het-challenge/. BAMSurgeon is available at: https://github.com/adamewing/bamsurgeon. The framework for subclonal mutation simulation is available at: http://search.cpan.org/~boutroslb/NGS-Tools-BAMSurgeon-v1.0.0/. The PhaseTools BAM phasing toolkit is available at https://github.com/mateidavid/phase-tools.

## Supplementary Note 1

### GISL

We developed a cascade ensemble model based on the Dirichlet process mixture model for tumour subclonal reconstruction. This model consists of four connected modules, named Module 1 (M1) to Module 4 (M4). Module M1 derives an initial estimate of cellularity from the phased phenotype information provided in the input Battenberg CNA data. This value is used in the subsequent steps to improve the accuracy of the decomposition results. Modules M2 and M3 predict the tumour subclonality based on a truncated Dirichlet process mixture model implemented by the blocked Gibbs sampler, but using different subsets of mutations. On the one hand, to consider the effect of CNAs, we perform the decomposition on a selected subset of mutations in M2 for which the total copy number exactly equals one. The expected variant allele frequencies for these mutation loci are deterministic and hence no arbitrary assumptions are needed. On the other hand, to reduce the effect of false positive mutation callings, we designed several filtering criteria making use of the information available in the input MuTect VCF data. Module M3 then performs the decomposition on the subset of mutations passing those filters. The above methods address the questions in SubChallenges 1 to 3. For SubChallenge 4, module M4 reconstructs the evolutionary relationships of the inferred subclones using a heuristic tree building method based on three assumptions: (a) infinite site (b) parsimony, and (c) that each subclone has no more than two child nodes.

This model has several advantages. First, the cascade ensemble architecture provides flexibility to customize for different practical application scenarios. Second, the core technique for inferring tumour subclones (modified truncated Dirichlet process mixture model) features the automatic generation of the number of components. Finally, during the subclonal reconstruction process, the method considers the effects from CNAs and the false positive mutation callings, and accordingly adjusts the predictions for improved accuracy.

### Object Integration

This method relies solely on the Battenberg copy-number profiles to re-estimate the purity. It first identifies segments that are not flagged as subclonal by Battenberg, and derives separate purity estimates from both the BAF and LogR. From the fitted integer values and the BAF and LogR, it is possible to back-calculate the purity for each segment. Object Integration summarizes these estimates using linear modeling and iterative fitting and the length of the segments as weights. It then takes a median between those independent purity estimates if the range is lower than 0.1 or takes the highest purity estimate if not.

