## Supplementary Figures for "Crowd-sourced benchmarking of single-sample tumour subclonal reconstruction"

### Supplementary Figure 1

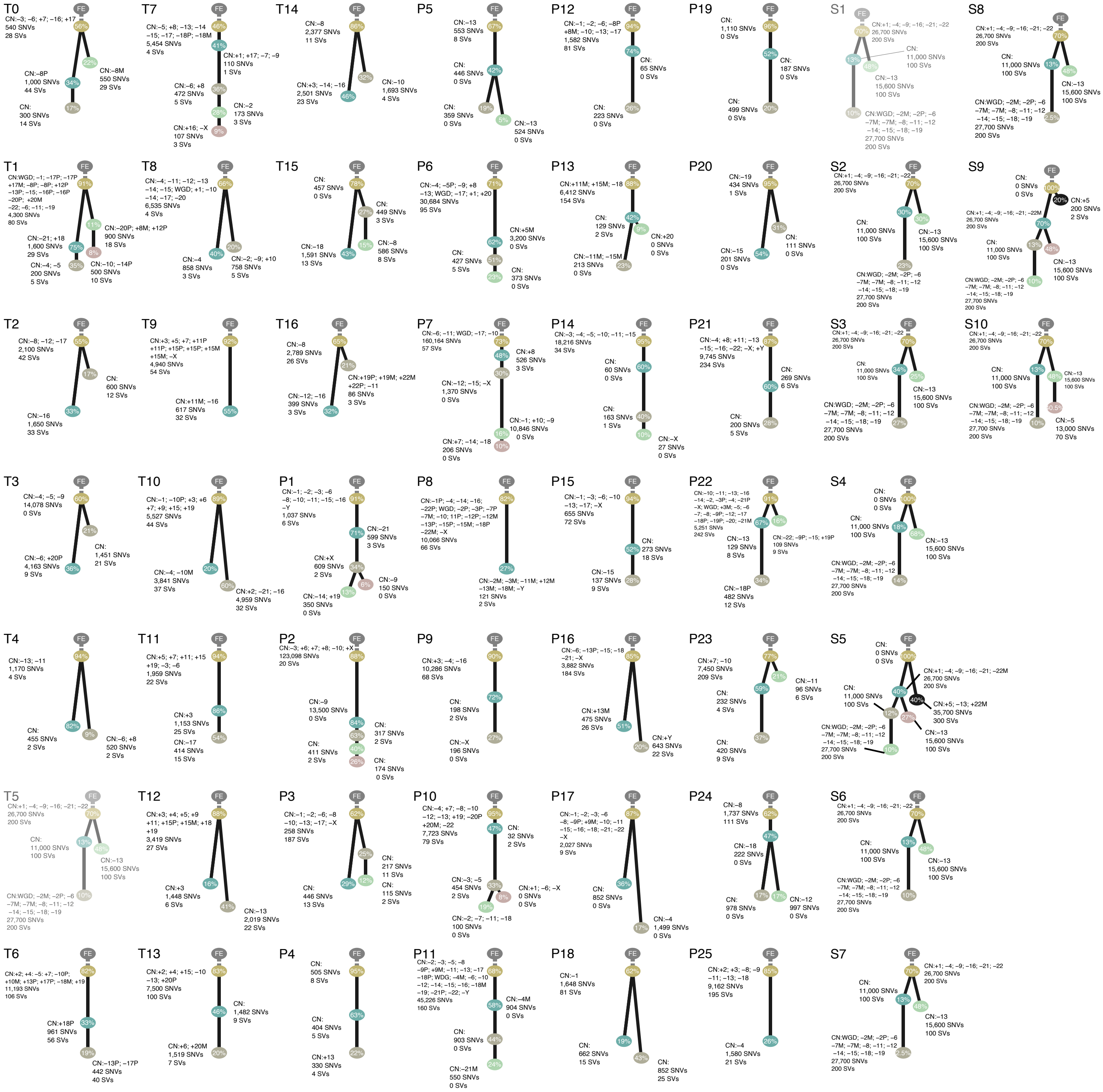

#### Supplementary Figure 2

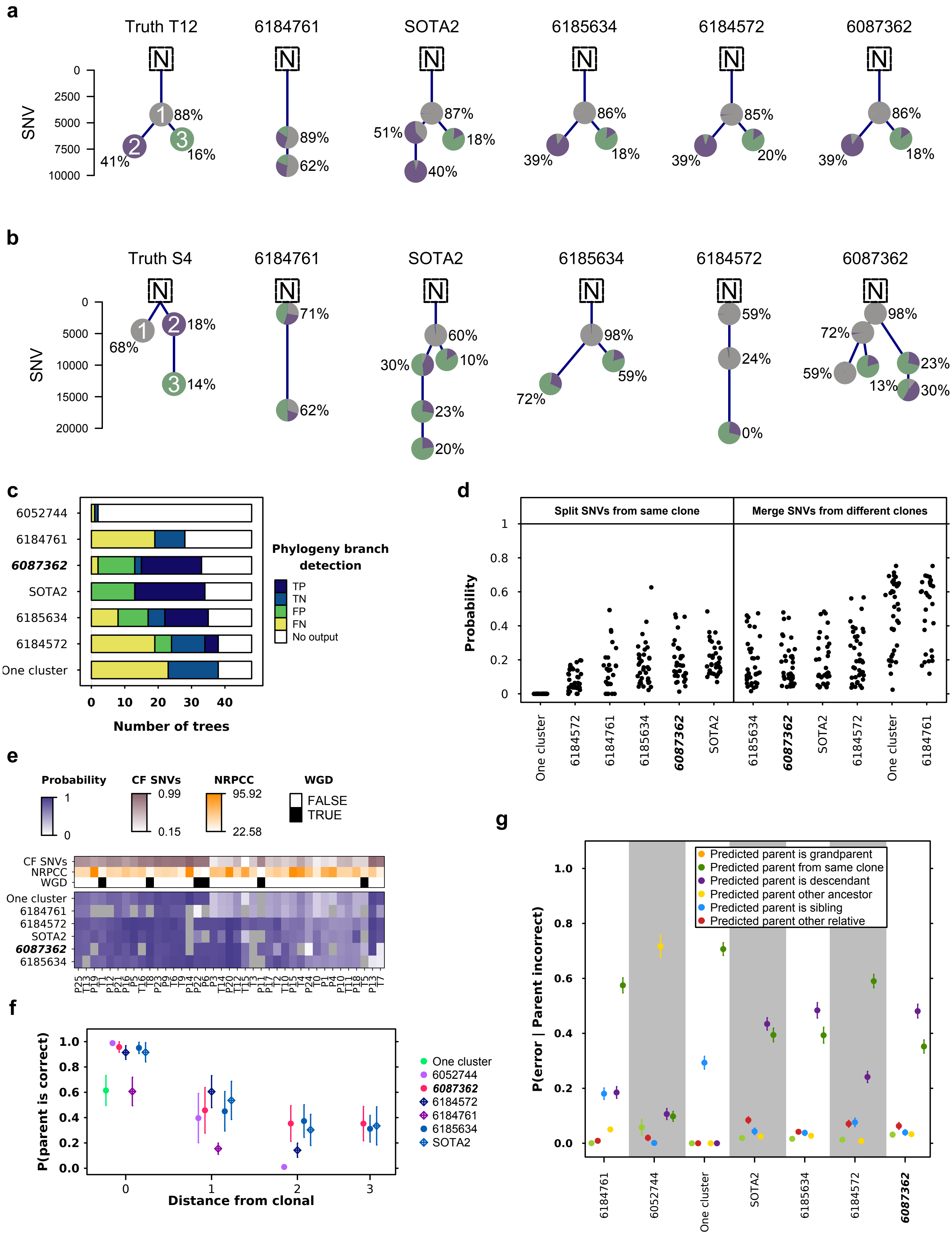

### Supplementary Figure 3

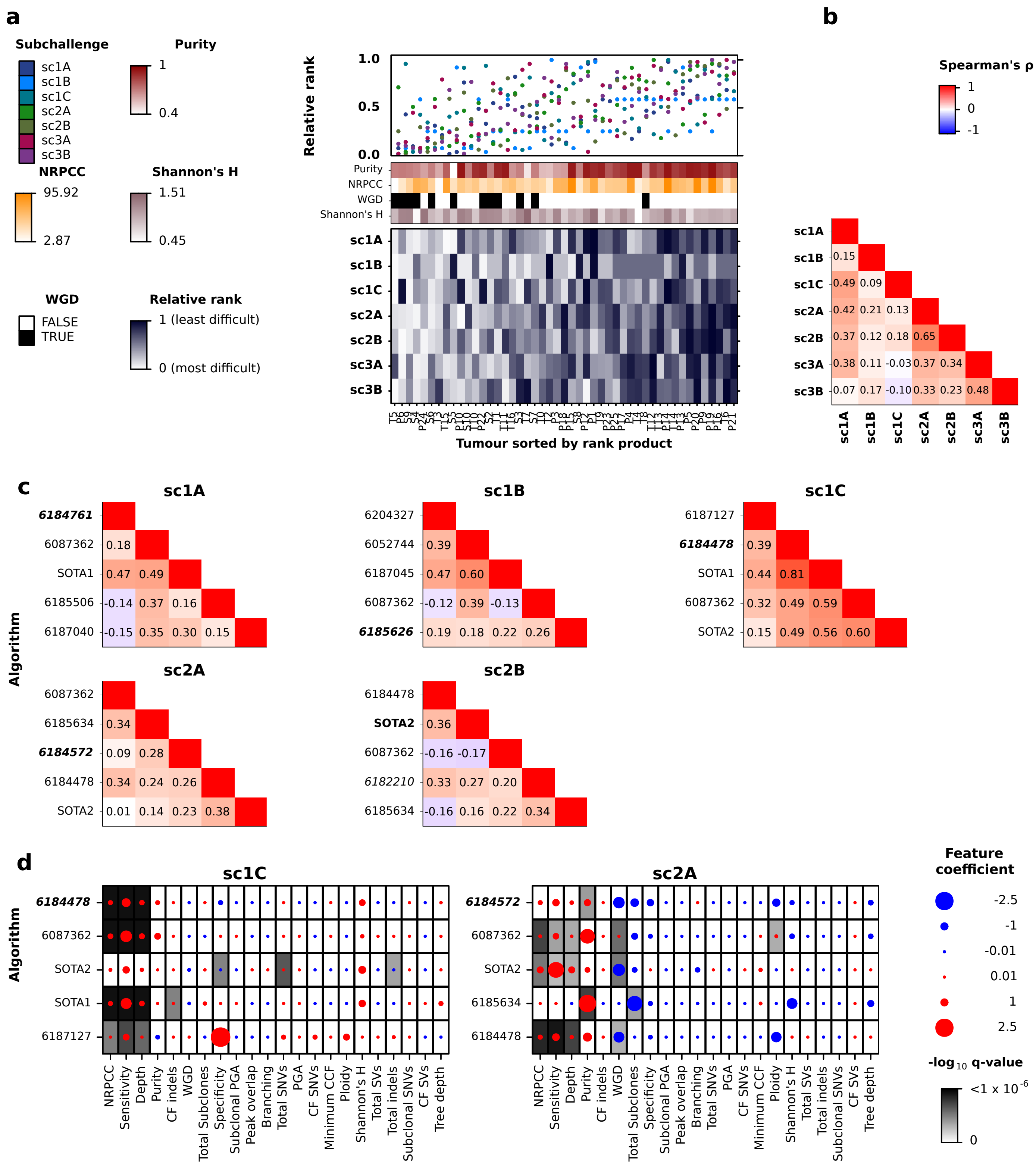

#### Supplementary Figure 4

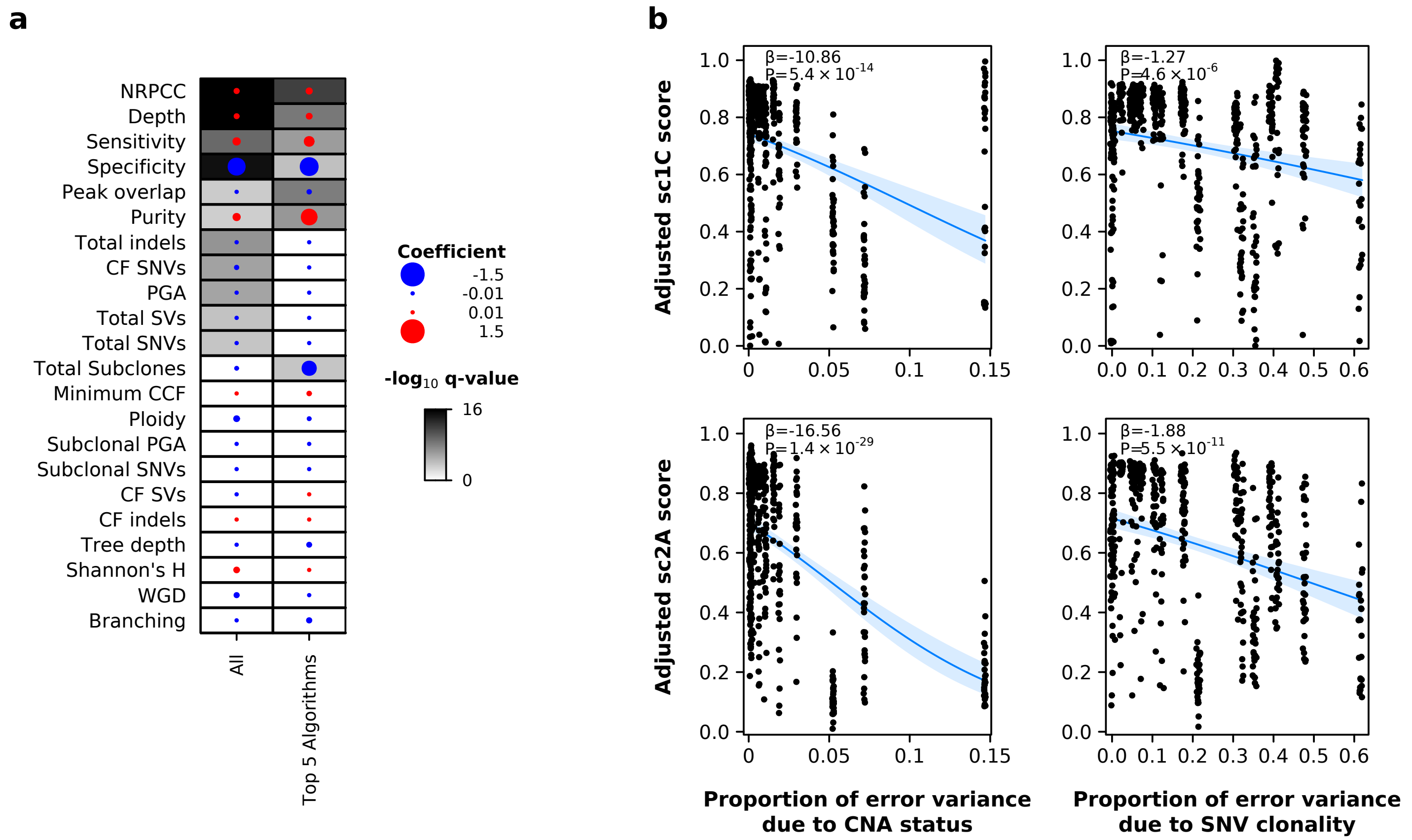

### Supplementary Figure 5

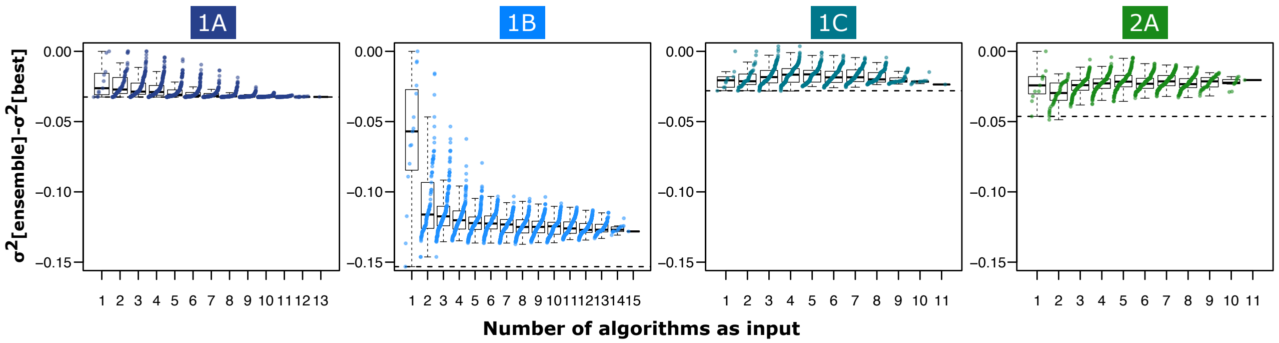
