## Supplementary Tables 4-6 for "Crowd-sourced benchmarking of single-sample tumour subclonal reconstruction"

**Supplementary Table 4: Regression results for clonal accuracy and SNV CP error.** $\beta$ regressions were used for accuracy and a linear model was used on inverse-normal transformed errors. N=4968 and N=4462484, respectively.

| | Estimate Accuracy | Std. Error Accuracy | P-value Accuracy | Estimate Error | Std. Error Error | P-value Error |
|---|---|---|---|---|---|---|
| (Intercept) | $-2.06\times10^{-1}$ | $1.48\times10^{-1}$ | $1.63\times10^{-1}$ | $5.26\times10^{-1}$ | $3.81\times10^{-3}$ | $< 2\times10^{-16}$ |
| clonal loss | $2.77\times10^{-2}$ | $7.59\times10^{-2}$ | $7.15\times10^{-1}$ | $1.98\times10^{-1}$ | $2.50\times10^{-3}$ | $< 2\times10^{-16}$ |
| clonal loss*SNV clonal | $2.60\times10^{-1}$ | $1.05\times10^{-1}$ | $1.29\times10^{-2}$ | $4.50\times10^{-2}$ | $2.80\times10^{-3}$ | $< 2\times10^{-16}$ |
| clonal gain | $1.21\times10^{-1}$ | $8.98\times10^{-2}$ | $1.76\times10^{-1}$ | $-1.53\times10^{-1}$ | $1.97\times10^{-3}$ | $< 2\times10^{-16}$ |
| clonal gain*SNV clonal | $-5.33\times10^{-2}$ | $1.21\times10^{-1}$ | $6.60\times10^{-1}$ | $9.99\times10^{-2}$ | $2.06\times10^{-3}$ | $< 2\times10^{-16}$ |
| subclonal loss | $6.55\times10^{-2}$ | $8.04\times10^{-2}$ | $4.15\times10^{-1}$ | $-2.13\times10^{-2}$ | $2.36\times10^{-3}$ | $< 2\times10^{-16}$ |
| subclonal loss*SNV clonal | $-3.29\times10^{-1}$ | $1.11\times10^{-1}$ | $3.06\times10^{-3}$ | $-1.43\times10^{-1}$ | $2.75\times10^{-3}$ | $< 2\times10^{-16}$ |
| subclonal gain | $-1.41\times10^{-1}$ | $8.91\times10^{-2}$ | $1.13\times10^{-1}$ | $2.09\times10^{-1}$ | $3.20\times10^{-3}$ | $< 2\times10^{-16}$ |
| subclonal gain*SNV clonal | $3.18\times10^{-1}$ | $1.20\times10^{-1}$ | $7.98\times10^{-3}$ | $-2.03\times10^{-1}$ | $3.63\times10^{-3}$ | $< 2\times10^{-16}$ |
| SNV clonal | $1.44\times10^{-1}$ | $7.28\times10^{-2}$ | $4.82\times10^{-2}$ | $-7.47\times10^{-1}$ | $1.16\times10^{-3}$ | $< 2\times10^{-16}$ |
| 6052744 | $5.68\times10^{-1}$ | $1.17\times10^{-1}$ | $1.21\times10^{-6}$ | -1.50 | $2.15\times10^{-3}$ | $< 2\times10^{-16}$ |
| 6087270 | $6.59\times10^{-1}$ | $1.15\times10^{-1}$ | $9.77\times10^{-9}$ | -1.37 | $2.06\times10^{-3}$ | $< 2\times10^{-16}$ |
| 6087309 | $6.92\times10^{-1}$ | $1.20\times10^{-1}$ | $6.97\times10^{-9}$ | $-4.88\times10^{-1}$ | $2.32\times10^{-3}$ | $< 2\times10^{-16}$ |
| 6087362 | $7.38\times10^{-1}$ | $1.15\times10^{-1}$ | $1.40\times10^{-10}$ | $-1.81\times10^{-1}$ | $2.05\times10^{-3}$ | $< 2\times10^{-16}$ |
| 6181022 | $4.68\times10^{-2}$ | $1.16\times10^{-1}$ | $6.86\times10^{-1}$ | $-9.63\times10^{-1}$ | $2.00\times10^{-3}$ | $< 2\times10^{-16}$ |
| 6184474 | $7.20\times10^{-1}$ | $1.14\times10^{-1}$ | $2.65\times10^{-10}$ | $-2.96\times10^{-1}$ | $2.00\times10^{-3}$ | $< 2\times10^{-16}$ |
| 6184478 | $7.24\times10^{-1}$ | $1.14\times10^{-1}$ | $2.10\times10^{-10}$ | $-2.36\times10^{-1}$ | $2.00\times10^{-3}$ | $< 2\times10^{-16}$ |
| 6184572 | $6.73\times10^{-1}$ | $1.15\times10^{-1}$ | $4.24\times10^{-9}$ | $-3.03\times10^{-1}$ | $2.04\times10^{-3}$ | $< 2\times10^{-16}$ |
| 6184761 | $9.40\times10^{-3}$ | $1.20\times10^{-1}$ | $9.38\times10^{-1}$ | $3.98\times10^{-1}$ | $2.19\times10^{-3}$ | $< 2\times10^{-16}$ |
| 6185634 | $7.97\times10^{-1}$ | $1.20\times10^{-1}$ | $3.02\times10^{-11}$ | $5.76\times10^{-2}$ | $2.31\times10^{-3}$ | $< 2\times10^{-16}$ |
| 6187040 | $7.49\times10^{-1}$ | $1.22\times10^{-1}$ | $9.65\times10^{-10}$ | $-1.63\times10^{-1}$ | $2.36\times10^{-3}$ | $< 2\times10^{-16}$ |
| 6187081 | $5.56\times10^{-1}$ | $1.19\times10^{-1}$ | $2.77\times10^{-6}$ | $-2.16\times10^{-1}$ | $2.11\times10^{-3}$ | $< 2\times10^{-16}$ |
| 6203106 | $5.00\times10^{-1}$ | $1.29\times10^{-1}$ | $1.06\times10^{-4}$ | $-4.05\times10^{-1}$ | $2.37\times10^{-3}$ | $< 2\times10^{-16}$ |
| 6400919 | $5.86\times10^{-1}$ | $1.15\times10^{-1}$ | $3.17\times10^{-7}$ | $-4.53\times10^{-1}$ | $2.00\times10^{-3}$ | $< 2\times10^{-16}$ |
| SOTA1 | $6.72\times10^{-1}$ | $1.15\times10^{-1}$ | $4.86\times10^{-9}$ | $-7.22\times10^{-2}$ | $2.06\times10^{-3}$ | $< 2\times10^{-16}$ |
| SOTA2 | $7.45\times10^{-1}$ | $1.15\times10^{-1}$ | $1.06\times10^{-10}$ | $-2.36\times10^{-1}$ | $2.19\times10^{-3}$ | $< 2\times10^{-16}$ |
| PCAWG-IR | $5.55\times10^{-1}$ | $1.15\times10^{-1}$ | $1.33\times10^{-6}$ | $-1.15\times10^{-1}$ | $2.00\times10^{-3}$ | $< 2\times10^{-16}$ |
| 6181088 | $7.41\times10^{-1}$ | $1.14\times10^{-1}$ | $7.48\times10^{-11}$ | $6.18\times10^{-1}$ | $2.00\times10^{-3}$ | $< 2\times10^{-16}$ |
| 6613462 | $8.28\times10^{-2}$ | $1.24\times10^{-1}$ | $5.03\times10^{-1}$ | $-6.63\times10^{-1}$ | $2.11\times10^{-3}$ | $< 2\times10^{-16}$ |
| P10 | $-1.28\times10^{-1}$ | $1.50\times10^{-1}$ | $3.94\times10^{-1}$ | $-1.82\times10^{-1}$ | $3.98\times10^{-3}$ | $< 2\times10^{-16}$ |
| P11 | $-2.24\times10^{-1}$ | $2.02\times10^{-1}$ | $2.68\times10^{-1}$ | $5.07\times10^{-1}$ | $3.89\times10^{-3}$ | $< 2\times10^{-16}$ |
| P12 | $2.35\times10^{-1}$ | $1.88\times10^{-1}$ | $2.11\times10^{-1}$ | $4.37\times10^{-2}$ | $5.36\times10^{-3}$ | $3.47\times10^{-16}$ |
| P13 | $4.92\times10^{-1}$ | $1.55\times10^{-1}$ | $1.52\times10^{-3}$ | $2.31\times10^{-1}$ | $4.14\times10^{-3}$ | $< 2\times10^{-16}$ |
| P14 | $1.75\times10^{-1}$ | $2.10\times10^{-1}$ | $4.04\times10^{-1}$ | $2.76\times10^{-1}$ | $3.84\times10^{-3}$ | $< 2\times10^{-16}$ |
| P15 | $3.81\times10^{-1}$ | $1.68\times10^{-1}$ | $2.34\times10^{-2}$ | $-6.73\times10^{-2}$ | $5.97\times10^{-3}$ | $< 2\times10^{-16}$ |
| P16 | $2.44\times10^{-1}$ | $1.67\times10^{-1}$ | $1.43\times10^{-1}$ | $1.90\times10^{-1}$ | $3.95\times10^{-3}$ | $< 2\times10^{-16}$ |
| P17 | $3.22\times10^{-1}$ | $1.64\times10^{-1}$ | $4.98\times10^{-2}$ | $1.11\times10^{-1}$ | $4.26\times10^{-3}$ | $< 2\times10^{-16}$ |
| P18 | $9.68\times10^{-3}$ | $1.91\times10^{-1}$ | $9.59\times10^{-1}$ | $5.78\times10^{-2}$ | $4.45\times10^{-3}$ | $< 2\times10^{-16}$ |
| P19 | $5.10\times10^{-1}$ | $2.59\times10^{-1}$ | $4.90\times10^{-2}$ | $2.02\times10^{-2}$ | $6.09\times10^{-3}$ | $9.16\times10^{-4}$ |
| P20 | $5.62\times10^{-1}$ | $1.62\times10^{-1}$ | $5.36\times10^{-4}$ | $-1.26\times10^{-1}$ | $6.16\times10^{-3}$ | $< 2\times10^{-16}$ |
| P21 | $7.89\times10^{-2}$ | $1.69\times10^{-1}$ | $6.41\times10^{-1}$ | $2.03\times10^{-1}$ | $3.90\times10^{-3}$ | $< 2\times10^{-16}$ |
| P22 | $-7.42\times10^{-4}$ | $1.55\times10^{-1}$ | $9.96\times10^{-1}$ | $1.77\times10^{-1}$ | $4.55\times10^{-3}$ | $< 2\times10^{-16}$ |
| P23 | $1.25\times10^{-2}$ | $1.56\times10^{-1}$ | $9.36\times10^{-1}$ | $2.30\times10^{-1}$ | $4.00\times10^{-3}$ | $< 2\times10^{-16}$ |
| P24 | $1.61\times10^{-1}$ | $1.73\times10^{-1}$ | $3.51\times10^{-1}$ | $1.66\times10^{-1}$ | $4.61\times10^{-3}$ | $< 2\times10^{-16}$ |
| P25 | $3.69\times10^{-1}$ | $1.57\times10^{-1}$ | $1.89\times10^{-2}$ | $2.40\times10^{-1}$ | $3.83\times10^{-3}$ | $< 2\times10^{-16}$ |
| P3 | $3.16\times10^{-1}$ | $1.84\times10^{-1}$ | $8.67\times10^{-2}$ | $-7.09\times10^{-2}$ | $6.28\times10^{-3}$ | $< 2\times10^{-16}$ |
| P4 | $3.06\times10^{-1}$ | $1.95\times10^{-1}$ | $1.16\times10^{-1}$ | $-1.96\times10^{-1}$ | $6.27\times10^{-3}$ | $< 2\times10^{-16}$ |
| P5 | $3.98\times10^{-1}$ | $1.66\times10^{-1}$ | $1.67\times10^{-2}$ | $6.49\times10^{-2}$ | $4.45\times10^{-3}$ | $< 2\times10^{-16}$ |
| P6 | $-1.09\times10^{-1}$ | $2.20\times10^{-1}$ | $6.20\times10^{-1}$ | $4.99\times10^{-1}$ | $3.88\times10^{-3}$ | $< 2\times10^{-16}$ |
| P8 | $2.49\times10^{-1}$ | $1.48\times10^{-1}$ | $9.29\times10^{-2}$ | $2.67\times10^{-1}$ | $4.06\times10^{-3}$ | $< 2\times10^{-16}$ |
| P9 | $2.85\times10^{-1}$ | $1.68\times10^{-1}$ | $8.91\times10^{-2}$ | $2.29\times10^{-1}$ | $3.88\times10^{-3}$ | $< 2\times10^{-16}$ |
| T0 | $-7.65\times10^{-2}$ | $1.56\times10^{-1}$ | $6.24\times10^{-1}$ | $-1.47\times10^{-1}$ | $5.00\times10^{-3}$ | $< 2\times10^{-16}$ |
| T1 | $9.67\times10^{-3}$ | $1.75\times10^{-1}$ | $9.56\times10^{-1}$ | $-1.21\times10^{-2}$ | $4.36\times10^{-3}$ | $5.56\times10^{-3}$ |
| T10 | $-2.58\times10^{-4}$ | $1.55\times10^{-1}$ | $9.99\times10^{-1}$ | $8.30\times10^{-2}$ | $3.86\times10^{-3}$ | $< 2\times10^{-16}$ |
| T11 | $-8.50\times10^{-2}$ | $1.48\times10^{-1}$ | $5.66\times10^{-1}$ | $-1.43\times10^{-1}$ | $4.52\times10^{-3}$ | $< 2\times10^{-16}$ |
| T12 | $2.98\times10^{-1}$ | $1.68\times10^{-1}$ | $7.66\times10^{-2}$ | $1.51\times10^{-1}$ | $4.14\times10^{-3}$ | $< 2\times10^{-16}$ |
| T13 | $1.36\times10^{-1}$ | $1.56\times10^{-1}$ | $3.86\times10^{-1}$ | $1.93\times10^{-1}$ | $3.91\times10^{-3}$ | $< 2\times10^{-16}$ |
| T14 | $1.17\times10^{-1}$ | $1.58\times10^{-1}$ | $4.57\times10^{-1}$ | $7.39\times10^{-2}$ | $4.14\times10^{-3}$ | $< 2\times10^{-16}$ |
| T15 | $1.78\times10^{-1}$ | $1.90\times10^{-1}$ | $3.50\times10^{-1}$ | $-1.44\times10^{-1}$ | $4.75\times10^{-3}$ | $< 2\times10^{-16}$ |
| T16 | $3.34\times10^{-1}$ | $1.54\times10^{-1}$ | $3.04\times10^{-2}$ | $3.67\times10^{-1}$ | $4.66\times10^{-3}$ | $< 2\times10^{-16}$ |
| T2 | $-1.47\times10^{-1}$ | $1.67\times10^{-1}$ | $3.80\times10^{-1}$ | $2.19\times10^{-1}$ | $4.43\times10^{-3}$ | $< 2\times10^{-16}$ |
| T3 | $-3.53\times10^{-1}$ | $1.60\times10^{-1}$ | $2.68\times10^{-2}$ | $5.55\times10^{-1}$ | $3.84\times10^{-3}$ | $< 2\times10^{-16}$ |
| T4 | $-1.41\times10^{-1}$ | $1.57\times10^{-1}$ | $3.69\times10^{-1}$ | $-5.23\times10^{-2}$ | $5.55\times10^{-3}$ | $< 2\times10^{-16}$ |
| T5 | $-4.01\times10^{-1}$ | $2.04\times10^{-1}$ | $4.94\times10^{-2}$ | $6.76\times10^{-1}$ | $5.02\times10^{-3}$ | $< 2\times10^{-16}$ |
| T6 | $9.12\times10^{-2}$ | $1.49\times10^{-1}$ | $5.40\times10^{-1}$ | $2.32\times10^{-1}$ | $3.88\times10^{-3}$ | $< 2\times10^{-16}$ |
| T7 | $-2.23\times10^{-1}$ | $1.49\times10^{-1}$ | $1.34\times10^{-1}$ | $1.51\times10^{-1}$ | $4.17\times10^{-3}$ | $< 2\times10^{-16}$ |
| T8 | $7.61\times10^{-2}$ | $1.57\times10^{-1}$ | $6.27\times10^{-1}$ | $5.57\times10^{-1}$ | $4.21\times10^{-3}$ | $< 2\times10^{-16}$ |
| T9 | $2.82\times10^{-2}$ | $1.58\times10^{-1}$ | $8.58\times10^{-1}$ | $-8.14\times10^{-3}$ | $4.21\times10^{-3}$ | $5.31\times10^{-2}$ |
| R-squared | $1.00\times10^{-1}$ | | | $4.10\times10^{-1}$ | | |
| Log-likelihood | $9.63\times10^{+3}$ | | | | | |
| phi | $5.44\times10^{-1}$ | $8.78\times10^{-3}$ | $< 2\times10^{-16}$ | | | |
| F-statistic | | | | $4.70\times10^{+4}$ | | |
| F-statistic DF | | | | $6.70\times10^{+1}$ | | |

**Supplementary Table 5: Regression results for clonal accuracy and SNV CP error.** $\beta$ regressions were used for accuracy and a linear model was used on inverse-normal transformed errors. N=4968 and N=4462484, respectively.

| | Estimate Accuracy | Std. Error Accuracy | P-value Accuracy | Estimate Error | Std. Error Error | P-value Error |
|---|---|---|---|---|---|---|
| (Intercept) | $-6.24 \times 10^{-2}$ | $1.48 \times 10^{-1}$ | $6.72 \times 10^{-1}$ | $-2.21 \times 10^{-1}$ | $3.81 \times 10^{-3}$ | $< 2 \times 10^{-16}$ |
| clonal loss | $2.88 \times 10^{-1}$ | $7.41 \times 10^{-2}$ | $1.03 \times 10^{-4}$ | $2.43 \times 10^{-1}$ | $1.40 \times 10^{-3}$ | $< 2 \times 10^{-16}$ |
| clonal loss*SNV subclonal | $-2.60 \times 10^{-1}$ | $1.05 \times 10^{-1}$ | $1.29 \times 10^{-2}$ | $-4.50 \times 10^{-2}$ | $2.80 \times 10^{-3}$ | $< 2 \times 10^{-16}$ |
| clonal gain | $6.81 \times 10^{-2}$ | $8.90 \times 10^{-2}$ | $4.44 \times 10^{-1}$ | $-5.29 \times 10^{-2}$ | $1.40 \times 10^{-3}$ | $< 2 \times 10^{-16}$ |
| clonal gain*SNV subclonal | $5.33 \times 10^{-2}$ | $1.21 \times 10^{-1}$ | $6.60 \times 10^{-1}$ | $-9.99 \times 10^{-2}$ | $2.06 \times 10^{-3}$ | $< 2 \times 10^{-16}$ |
| subclonal loss | $-2.63 \times 10^{-1}$ | $8.06 \times 10^{-2}$ | $1.09 \times 10^{-3}$ | $-1.64 \times 10^{-1}$ | $1.69 \times 10^{-3}$ | $< 2 \times 10^{-16}$ |
| subclonal loss*SNV subclonal | $3.29 \times 10^{-1}$ | $1.11 \times 10^{-1}$ | $3.06 \times 10^{-3}$ | $1.43 \times 10^{-1}$ | $2.75 \times 10^{-3}$ | $< 2 \times 10^{-16}$ |
| subclonal gain | $1.77 \times 10^{-1}$ | $8.71 \times 10^{-2}$ | $4.21 \times 10^{-2}$ | $6.24 \times 10^{-3}$ | $2.11 \times 10^{-3}$ | $3.11 \times 10^{-3}$ |
| subclonal gain*SNV subclonal | $-3.18 \times 10^{-1}$ | $1.20 \times 10^{-1}$ | $7.98 \times 10^{-3}$ | $2.03 \times 10^{-1}$ | $3.63 \times 10^{-3}$ | $< 2 \times 10^{-16}$ |
| SNV subclonal | $-1.44 \times 10^{-1}$ | $7.28 \times 10^{-2}$ | $4.82 \times 10^{-2}$ | $7.47 \times 10^{-1}$ | $1.16 \times 10^{-3}$ | $< 2 \times 10^{-16}$ |
| 6052744 | $5.68 \times 10^{-1}$ | $1.17 \times 10^{-1}$ | $1.21 \times 10^{-6}$ | -1.50 | $2.15 \times 10^{-3}$ | $< 2 \times 10^{-16}$ |
| 6087270 | $6.59 \times 10^{-1}$ | $1.15 \times 10^{-1}$ | $9.77 \times 10^{-9}$ | -1.37 | $2.06 \times 10^{-3}$ | $< 2 \times 10^{-16}$ |
| 6087309 | $6.92 \times 10^{-1}$ | $1.20 \times 10^{-1}$ | $6.97 \times 10^{-9}$ | $-4.88 \times 10^{-1}$ | $2.32 \times 10^{-3}$ | $< 2 \times 10^{-16}$ |
| 6087362 | $7.38 \times 10^{-1}$ | $1.15 \times 10^{-1}$ | $1.40 \times 10^{-10}$ | $-1.81 \times 10^{-1}$ | $2.05 \times 10^{-3}$ | $< 2 \times 10^{-16}$ |
| 6181022 | $4.68 \times 10^{-2}$ | $1.16 \times 10^{-1}$ | $6.86 \times 10^{-1}$ | $-9.63 \times 10^{-1}$ | $2.00 \times 10^{-3}$ | $< 2 \times 10^{-16}$ |
| 6184474 | $7.20 \times 10^{-1}$ | $1.14 \times 10^{-1}$ | $2.65 \times 10^{-10}$ | $-2.96 \times 10^{-1}$ | $2.00 \times 10^{-3}$ | $< 2 \times 10^{-16}$ |
| 6184478 | $7.24 \times 10^{-1}$ | $1.14 \times 10^{-1}$ | $2.10 \times 10^{-10}$ | $-2.36 \times 10^{-1}$ | $2.00 \times 10^{-3}$ | $< 2 \times 10^{-16}$ |
| 6184572 | $6.73 \times 10^{-1}$ | $1.15 \times 10^{-1}$ | $4.24 \times 10^{-9}$ | $-3.03 \times 10^{-1}$ | $2.04 \times 10^{-3}$ | $< 2 \times 10^{-16}$ |
| 6184761 | $9.40 \times 10^{-3}$ | $1.20 \times 10^{-1}$ | $9.38 \times 10^{-1}$ | $3.98 \times 10^{-1}$ | $2.19 \times 10^{-3}$ | $< 2 \times 10^{-16}$ |
| 6185634 | $7.97 \times 10^{-1}$ | $1.20 \times 10^{-1}$ | $3.02 \times 10^{-11}$ | $5.76 \times 10^{-2}$ | $2.31 \times 10^{-3}$ | $< 2 \times 10^{-16}$ |
| 6187040 | $7.49 \times 10^{-1}$ | $1.22 \times 10^{-1}$ | $9.65 \times 10^{-10}$ | $-1.63 \times 10^{-1}$ | $2.36 \times 10^{-3}$ | $< 2 \times 10^{-16}$ |
| 6187081 | $5.56 \times 10^{-1}$ | $1.19 \times 10^{-1}$ | $2.77 \times 10^{-6}$ | $-2.16 \times 10^{-1}$ | $2.11 \times 10^{-3}$ | $< 2 \times 10^{-16}$ |
| 6203106 | $5.00 \times 10^{-1}$ | $1.29 \times 10^{-1}$ | $1.06 \times 10^{-4}$ | $-4.05 \times 10^{-1}$ | $2.37 \times 10^{-3}$ | $< 2 \times 10^{-16}$ |
| 6400919 | $5.86 \times 10^{-1}$ | $1.15 \times 10^{-1}$ | $3.17 \times 10^{-7}$ | $-4.53 \times 10^{-1}$ | $2.00 \times 10^{-3}$ | $< 2 \times 10^{-16}$ |
| SOTA1 | $6.72 \times 10^{-1}$ | $1.15 \times 10^{-1}$ | $4.86 \times 10^{-9}$ | $-7.22 \times 10^{-2}$ | $2.06 \times 10^{-3}$ | $< 2 \times 10^{-16}$ |
| SOTA2 | $7.45 \times 10^{-1}$ | $1.15 \times 10^{-1}$ | $1.06 \times 10^{-10}$ | $-2.36 \times 10^{-1}$ | $2.19 \times 10^{-3}$ | $< 2 \times 10^{-16}$ |
| PCAWG-IR | $5.55 \times 10^{-1}$ | $1.15 \times 10^{-1}$ | $1.33 \times 10^{-6}$ | $-1.15 \times 10^{-1}$ | $2.00 \times 10^{-3}$ | $< 2 \times 10^{-16}$ |
| 6181088 | $7.41 \times 10^{-1}$ | $1.14 \times 10^{-1}$ | $7.48 \times 10^{-11}$ | $6.18 \times 10^{-1}$ | $2.00 \times 10^{-3}$ | $< 2 \times 10^{-16}$ |
| 6613462 | $8.28 \times 10^{-2}$ | $1.24 \times 10^{-1}$ | $5.03 \times 10^{-1}$ | $-6.63 \times 10^{-1}$ | $2.11 \times 10^{-3}$ | $< 2 \times 10^{-16}$ |
| P10 | $-1.28 \times 10^{-1}$ | $1.50 \times 10^{-1}$ | $3.94 \times 10^{-1}$ | $-1.82 \times 10^{-1}$ | $3.98 \times 10^{-3}$ | $< 2 \times 10^{-16}$ |
| P11 | $-2.24 \times 10^{-1}$ | $2.02 \times 10^{-1}$ | $2.68 \times 10^{-1}$ | $5.07 \times 10^{-1}$ | $3.89 \times 10^{-3}$ | $< 2 \times 10^{-16}$ |
| P12 | $2.35 \times 10^{-1}$ | $1.88 \times 10^{-1}$ | $2.11 \times 10^{-1}$ | $4.37 \times 10^{-2}$ | $5.36 \times 10^{-3}$ | $3.47 \times 10^{-16}$ |
| P13 | $4.92 \times 10^{-1}$ | $1.55 \times 10^{-1}$ | $1.52 \times 10^{-3}$ | $2.31 \times 10^{-1}$ | $4.14 \times 10^{-3}$ | $< 2 \times 10^{-16}$ |
| P14 | $1.75 \times 10^{-1}$ | $2.10 \times 10^{-1}$ | $4.04 \times 10^{-1}$ | $2.76 \times 10^{-1}$ | $3.84 \times 10^{-3}$ | $< 2 \times 10^{-16}$ |
| P15 | $3.81 \times 10^{-1}$ | $1.68 \times 10^{-1}$ | $2.34 \times 10^{-2}$ | $-6.73 \times 10^{-2}$ | $5.97 \times 10^{-3}$ | $< 2 \times 10^{-16}$ |
| P16 | $2.44 \times 10^{-1}$ | $1.67 \times 10^{-1}$ | $1.43 \times 10^{-1}$ | $1.90 \times 10^{-1}$ | $3.95 \times 10^{-3}$ | $< 2 \times 10^{-16}$ |
| P17 | $3.22 \times 10^{-1}$ | $1.64 \times 10^{-1}$ | $4.98 \times 10^{-2}$ | $1.11 \times 10^{-1}$ | $4.26 \times 10^{-3}$ | $< 2 \times 10^{-16}$ |
| P18 | $9.68 \times 10^{-3}$ | $1.91 \times 10^{-1}$ | $9.59 \times 10^{-1}$ | $5.78 \times 10^{-2}$ | $4.45 \times 10^{-3}$ | $< 2 \times 10^{-16}$ |
| P19 | $5.10 \times 10^{-1}$ | $2.59 \times 10^{-1}$ | $4.90 \times 10^{-2}$ | $2.02 \times 10^{-2}$ | $6.09 \times 10^{-3}$ | $9.16 \times 10^{-4}$ |
| P20 | $5.62 \times 10^{-1}$ | $1.62 \times 10^{-1}$ | $5.36 \times 10^{-4}$ | $-1.26 \times 10^{-1}$ | $6.16 \times 10^{-3}$ | $< 2 \times 10^{-16}$ |
| P21 | $7.89 \times 10^{-2}$ | $1.69 \times 10^{-1}$ | $6.41 \times 10^{-1}$ | $2.03 \times 10^{-1}$ | $3.90 \times 10^{-3}$ | $< 2 \times 10^{-16}$ |
| P22 | $-7.42 \times 10^{-4}$ | $1.55 \times 10^{-1}$ | $9.96 \times 10^{-1}$ | $1.77 \times 10^{-1}$ | $4.55 \times 10^{-3}$ | $< 2 \times 10^{-16}$ |
| P23 | $1.25 \times 10^{-2}$ | $1.56 \times 10^{-1}$ | $9.36 \times 10^{-1}$ | $2.30 \times 10^{-1}$ | $4.00 \times 10^{-3}$ | $< 2 \times 10^{-16}$ |
| P24 | $1.61 \times 10^{-1}$ | $1.73 \times 10^{-1}$ | $3.51 \times 10^{-1}$ | $1.66 \times 10^{-1}$ | $4.61 \times 10^{-3}$ | $< 2 \times 10^{-16}$ |
| P25 | $3.69 \times 10^{-1}$ | $1.57 \times 10^{-1}$ | $1.89 \times 10^{-2}$ | $2.40 \times 10^{-1}$ | $3.83 \times 10^{-3}$ | $< 2 \times 10^{-16}$ |
| P3 | $3.16 \times 10^{-1}$ | $1.84 \times 10^{-1}$ | $8.67 \times 10^{-2}$ | $-7.09 \times 10^{-2}$ | $6.28 \times 10^{-3}$ | $< 2 \times 10^{-16}$ |
| P4 | $3.06 \times 10^{-1}$ | $1.95 \times 10^{-1}$ | $1.16 \times 10^{-1}$ | $-1.96 \times 10^{-1}$ | $6.27 \times 10^{-3}$ | $< 2 \times 10^{-16}$ |
| P5 | $3.98 \times 10^{-1}$ | $1.66 \times 10^{-1}$ | $1.67 \times 10^{-2}$ | $6.49 \times 10^{-2}$ | $4.45 \times 10^{-3}$ | $< 2 \times 10^{-16}$ |
| P6 | $-1.09 \times 10^{-1}$ | $2.20 \times 10^{-1}$ | $6.20 \times 10^{-1}$ | $4.99 \times 10^{-1}$ | $3.88 \times 10^{-3}$ | $< 2 \times 10^{-16}$ |
| P8 | $2.49 \times 10^{-1}$ | $1.48 \times 10^{-1}$ | $9.29 \times 10^{-2}$ | $2.67 \times 10^{-1}$ | $4.06 \times 10^{-3}$ | $< 2 \times 10^{-16}$ |
| P9 | $2.85 \times 10^{-1}$ | $1.68 \times 10^{-1}$ | $8.91 \times 10^{-2}$ | $2.29 \times 10^{-1}$ | $3.88 \times 10^{-3}$ | $< 2 \times 10^{-16}$ |
| T0 | $-7.65 \times 10^{-2}$ | $1.56 \times 10^{-1}$ | $6.24 \times 10^{-1}$ | $-1.47 \times 10^{-1}$ | $5.00 \times 10^{-3}$ | $< 2 \times 10^{-16}$ |
| T1 | $9.67 \times 10^{-3}$ | $1.75 \times 10^{-1}$ | $9.56 \times 10^{-1}$ | $-1.21 \times 10^{-2}$ | $4.36 \times 10^{-3}$ | $5.56 \times 10^{-3}$ |
| T10 | $-2.58 \times 10^{-4}$ | $1.55 \times 10^{-1}$ | $9.99 \times 10^{-1}$ | $8.30 \times 10^{-2}$ | $3.86 \times 10^{-3}$ | $< 2 \times 10^{-16}$ |
| T11 | $-8.50 \times 10^{-2}$ | $1.48 \times 10^{-1}$ | $5.66 \times 10^{-1}$ | $-1.43 \times 10^{-1}$ | $4.52 \times 10^{-3}$ | $< 2 \times 10^{-16}$ |
| T12 | $2.98 \times 10^{-1}$ | $1.68 \times 10^{-1}$ | $7.66 \times 10^{-2}$ | $1.51 \times 10^{-1}$ | $4.14 \times 10^{-3}$ | $< 2 \times 10^{-16}$ |
| T13 | $1.36 \times 10^{-1}$ | $1.56 \times 10^{-1}$ | $3.86 \times 10^{-1}$ | $1.93 \times 10^{-1}$ | $3.91 \times 10^{-3}$ | $< 2 \times 10^{-16}$ |
| T14 | $1.17 \times 10^{-1}$ | $1.58 \times 10^{-1}$ | $4.57 \times 10^{-1}$ | $7.39 \times 10^{-2}$ | $4.14 \times 10^{-3}$ | $< 2 \times 10^{-16}$ |
| T15 | $1.78 \times 10^{-1}$ | $1.90 \times 10^{-1}$ | $3.50 \times 10^{-1}$ | $-1.44 \times 10^{-1}$ | $4.75 \times 10^{-3}$ | $< 2 \times 10^{-16}$ |
| T16 | $3.34 \times 10^{-1}$ | $1.54 \times 10^{-1}$ | $3.04 \times 10^{-2}$ | $3.67 \times 10^{-1}$ | $4.66 \times 10^{-3}$ | $< 2 \times 10^{-16}$ |
| T2 | $-1.47 \times 10^{-1}$ | $1.67 \times 10^{-1}$ | $3.80 \times 10^{-1}$ | $2.19 \times 10^{-1}$ | $4.43 \times 10^{-3}$ | $< 2 \times 10^{-16}$ |
| T3 | $-3.53 \times 10^{-1}$ | $1.60 \times 10^{-1}$ | $2.68 \times 10^{-2}$ | $5.55 \times 10^{-1}$ | $3.84 \times 10^{-3}$ | $< 2 \times 10^{-16}$ |
| T4 | $-1.41 \times 10^{-1}$ | $1.57 \times 10^{-1}$ | $3.69 \times 10^{-1}$ | $-5.23 \times 10^{-2}$ | $5.55 \times 10^{-3}$ | $< 2 \times 10^{-16}$ |
| T5 | $-4.01 \times 10^{-1}$ | $2.04 \times 10^{-1}$ | $4.94 \times 10^{-2}$ | $6.76 \times 10^{-1}$ | $5.02 \times 10^{-3}$ | $< 2 \times 10^{-16}$ |
| T6 | $9.12 \times 10^{-2}$ | $1.49 \times 10^{-1}$ | $5.40 \times 10^{-1}$ | $2.32 \times 10^{-1}$ | $3.88 \times 10^{-3}$ | $< 2 \times 10^{-16}$ |
| T7 | $-2.23 \times 10^{-1}$ | $1.49 \times 10^{-1}$ | $1.34 \times 10^{-1}$ | $1.51 \times 10^{-1}$ | $4.17 \times 10^{-3}$ | $< 2 \times 10^{-16}$ |
| T8 | $7.61 \times 10^{-2}$ | $1.57 \times 10^{-1}$ | $6.27 \times 10^{-1}$ | $5.57 \times 10^{-1}$ | $4.21 \times 10^{-3}$ | $< 2 \times 10^{-16}$ |
| T9 | $2.82 \times 10^{-2}$ | $1.58 \times 10^{-1}$ | $8.58 \times 10^{-1}$ | $-8.14 \times 10^{-3}$ | $4.21 \times 10^{-3}$ | $5.31 \times 10^{-2}$ |
| R-squared | $1.00 \times 10^{-1}$ | | | $4.10 \times 10^{-1}$ | | |
| Log-likelihood | $9.63 \times 10^{+3}$ | | | | | |
| phi | $5.44 \times 10^{-1}$ | $8.78 \times 10^{-3}$ | $< 2 \times 10^{-16}$ | | | |
| F-statistic | | | | $4.70 \times 10^{+4}$ | | |
| F-statistic DF | | | | $6.70 \times 10^{+1}$ | | |

**Supplementary Table 6: Regression results assessing the effect of Battenberg CNA accuracy clonal accuracy and SNV CP error in subclonal CNAs.** $\beta$ regressions were used for accuracy and a linear model was used on inverse-normal transformed errors. N=5212 and N=4462484, respectively.

| | Estimate Accuracy | Std. Error Accuracy | P-value Accuracy | Estimate Error | Std. Error Error | P-value Error |
|---|---|---|---|---|---|---|
| (Intercept) | -1.91x10$^{-1}$ | 1.39x10$^{-1}$ | 1.69x10$^{-1}$ | 6.77x10$^{-1}$ | 4.04x10$^{-3}$ | $<$ 2x10$^{-16}$ |
| Battenberg incorrect | 6.02x10$^{-2}$ | 7.78x10$^{-2}$ | 4.39x10$^{-1}$ | 1.12x10$^{-1}$ | 2.73x10$^{-3}$ | $<$ 2x10$^{-16}$ |
| clonal loss | 2.75x10$^{-2}$ | 7.61x10$^{-2}$ | 7.17x10$^{-1}$ | 2.12x10$^{-1}$ | 2.65x10$^{-3}$ | $<$ 2x10$^{-16}$ |
| clonal loss*SNV clonal | 2.55x10$^{-1}$ | 1.05x10$^{-1}$ | 1.49x10$^{-2}$ | 4.27x10$^{-2}$ | 2.97x10$^{-3}$ | $<$ 2x10$^{-16}$ |
| clonal gain | 1.24x10$^{-1}$ | 9.00x10$^{-2}$ | 1.69x10$^{-1}$ | -1.52x10$^{-1}$ | 2.09x10$^{-3}$ | $<$ 2x10$^{-16}$ |
| clonal gain*SNV clonal | -5.26x10$^{-2}$ | 1.21x10$^{-1}$ | 6.65x10$^{-1}$ | 1.01x10$^{-1}$ | 2.18x10$^{-3}$ | $<$ 2x10$^{-16}$ |
| subclonal loss | 2.99x10$^{-2}$ | 7.87x10$^{-2}$ | 7.05x10$^{-1}$ | -2.74x10$^{-2}$ | 2.50x10$^{-3}$ | $<$ 2x10$^{-16}$ |
| subclonal loss*SNV clonal | -2.59x10$^{-1}$ | 1.08x10$^{-1}$ | 1.58x10$^{-2}$ | -1.53x10$^{-1}$ | 2.92x10$^{-3}$ | $<$ 2x10$^{-16}$ |
| subclonal gain | -2.25x10$^{-1}$ | 8.85x10$^{-2}$ | 1.12x10$^{-2}$ | 1.85x10$^{-1}$ | 3.58x10$^{-3}$ | $<$ 2x10$^{-16}$ |
| subclonal gain*SNV clonal | 4.56x10$^{-1}$ | 1.15x10$^{-1}$ | 7.75x10$^{-5}$ | -2.33x10$^{-1}$ | 3.85x10$^{-3}$ | $<$ 2x10$^{-16}$ |
| SNV clonal | 1.40x10$^{-1}$ | 7.30x10$^{-2}$ | 5.43x10$^{-2}$ | -8.00x10$^{-1}$ | 1.23x10$^{-3}$ | $<$ 2x10$^{-16}$ |
| 6052744 | 5.38x10$^{-1}$ | 1.14x10$^{-1}$ | 2.53x10$^{-6}$ | -1.58 | 2.28x10$^{-3}$ | $<$ 2x10$^{-16}$ |
| 6087270 | 6.33x10$^{-1}$ | 1.12x10$^{-1}$ | 1.71x10$^{-8}$ | -1.43 | 2.19x10$^{-3}$ | $<$ 2x10$^{-16}$ |
| 6087309 | 6.62x10$^{-1}$ | 1.17x10$^{-1}$ | 1.53x10$^{-8}$ | -5.29x10$^{-1}$ | 2.46x10$^{-3}$ | $<$ 2x10$^{-16}$ |
| 6087362 | 7.01x10$^{-1}$ | 1.12x10$^{-1}$ | 4.50x10$^{-10}$ | -2.02x10$^{-1}$ | 2.17x10$^{-3}$ | $<$ 2x10$^{-16}$ |
| 6181022 | 4.25x10$^{-2}$ | 1.13x10$^{-1}$ | 7.07x10$^{-1}$ | -1.01 | 2.12x10$^{-3}$ | $<$ 2x10$^{-16}$ |
| 6184474 | 6.81x10$^{-1}$ | 1.11x10$^{-1}$ | 9.60x10$^{-10}$ | -3.30x10$^{-1}$ | 2.12x10$^{-3}$ | $<$ 2x10$^{-16}$ |
| 6184478 | 6.86x10$^{-1}$ | 1.11x10$^{-1}$ | 7.28x10$^{-10}$ | -2.67x10$^{-1}$ | 2.12x10$^{-3}$ | $<$ 2x10$^{-16}$ |
| 6184572 | 6.33x10$^{-1}$ | 1.12x10$^{-1}$ | 1.56x10$^{-8}$ | -3.36x10$^{-1}$ | 2.16x10$^{-3}$ | $<$ 2x10$^{-16}$ |
| 6184761 | -2.57x10$^{-4}$ | 1.17x10$^{-1}$ | 9.98x10$^{-1}$ | 4.32x10$^{-1}$ | 2.32x10$^{-3}$ | $<$ 2x10$^{-16}$ |
| 6185634 | 7.68x10$^{-1}$ | 1.18x10$^{-1}$ | 8.41x10$^{-11}$ | 4.85x10$^{-2}$ | 2.44x10$^{-3}$ | $<$ 2x10$^{-16}$ |
| 6187040 | 7.14x10$^{-1}$ | 1.20x10$^{-1}$ | 2.81x10$^{-9}$ | -1.76x10$^{-1}$ | 2.50x10$^{-3}$ | $<$ 2x10$^{-16}$ |
| 6187081 | 5.25x10$^{-1}$ | 1.16x10$^{-1}$ | 5.74x10$^{-6}$ | -2.41x10$^{-1}$ | 2.23x10$^{-3}$ | $<$ 2x10$^{-16}$ |
| 6203106 | 4.66x10$^{-1}$ | 1.27x10$^{-1}$ | 2.42x10$^{-4}$ | -4.41x10$^{-1}$ | 2.52x10$^{-3}$ | $<$ 2x10$^{-16}$ |
| 6400919 | 5.49x10$^{-1}$ | 1.12x10$^{-1}$ | 9.34x10$^{-7}$ | -4.94x10$^{-1}$ | 2.12x10$^{-3}$ | $<$ 2x10$^{-16}$ |
| SOTA1 | 6.35x10$^{-1}$ | 1.12x10$^{-1}$ | 1.50x10$^{-8}$ | -8.84x10$^{-2}$ | 2.19x10$^{-3}$ | $<$ 2x10$^{-16}$ |
| SOTA2 | 7.06x10$^{-1}$ | 1.13x10$^{-1}$ | 3.75x10$^{-10}$ | -2.63x10$^{-1}$ | 2.32x10$^{-3}$ | $<$ 2x10$^{-16}$ |
| PCAWG-IR | 5.28x10$^{-1}$ | 1.12x10$^{-1}$ | 2.50x10$^{-6}$ | -1.32x10$^{-1}$ | 2.12x10$^{-3}$ | $<$ 2x10$^{-16}$ |
| 6181088 | 7.04x10$^{-1}$ | 1.11x10$^{-1}$ | 2.47x10$^{-10}$ | 6.53x10$^{-1}$ | 2.12x10$^{-3}$ | $<$ 2x10$^{-16}$ |
| 6613462 | 8.12x10$^{-2}$ | 1.21x10$^{-1}$ | 5.01x10$^{-1}$ | -7.08x10$^{-1}$ | 2.23x10$^{-3}$ | $<$ 2x10$^{-16}$ |
| P10 | -1.28x10$^{-1}$ | 1.42x10$^{-1}$ | 3.67x10$^{-1}$ | -1.93x10$^{-1}$ | 4.22x10$^{-3}$ | $<$ 2x10$^{-16}$ |
| P11 | -2.14x10$^{-1}$ | 1.96x10$^{-1}$ | 2.75x10$^{-1}$ | 5.25x10$^{-1}$ | 4.13x10$^{-3}$ | $<$ 2x10$^{-16}$ |
| P12 | 2.39x10$^{-1}$ | 1.82x10$^{-1}$ | 1.91x10$^{-1}$ | 5.19x10$^{-2}$ | 5.68x10$^{-3}$ | $<$ 2x10$^{-16}$ |
| P13 | 5.05x10$^{-1}$ | 1.48x10$^{-1}$ | 6.48x10$^{-4}$ | 2.42x10$^{-1}$ | 4.40x10$^{-3}$ | $<$ 2x10$^{-16}$ |
| P14 | 1.78x10$^{-1}$ | 2.05x10$^{-1}$ | 3.85x10$^{-1}$ | 2.94x10$^{-1}$ | 4.07x10$^{-3}$ | $<$ 2x10$^{-16}$ |
| P15 | 3.88x10$^{-1}$ | 1.61x10$^{-1}$ | 1.61x10$^{-2}$ | -6.72x10$^{-2}$ | 6.33x10$^{-3}$ | $<$ 2x10$^{-16}$ |
| P16 | 2.50x10$^{-1}$ | 1.61x10$^{-1}$ | 1.20x10$^{-1}$ | 2.00x10$^{-1}$ | 4.19x10$^{-3}$ | $<$ 2x10$^{-16}$ |
| P17 | 3.27x10$^{-1}$ | 1.57x10$^{-1}$ | 3.75x10$^{-2}$ | 1.16x10$^{-1}$ | 4.52x10$^{-3}$ | $<$ 2x10$^{-16}$ |
| P18 | 1.59x10$^{-2}$ | 1.85x10$^{-1}$ | 9.31x10$^{-1}$ | 5.02x10$^{-2}$ | 4.72x10$^{-3}$ | $<$ 2x10$^{-16}$ |
| P19 | 5.11x10$^{-1}$ | 2.55x10$^{-1}$ | 4.50x10$^{-2}$ | 1.94x10$^{-2}$ | 6.46x10$^{-3}$ | 2.66x10$^{-3}$ |
| P20 | 5.66x10$^{-1}$ | 1.55x10$^{-1}$ | 2.54x10$^{-4}$ | -1.27x10$^{-1}$ | 6.53x10$^{-3}$ | $<$ 2x10$^{-16}$ |
| P21 | 8.29x10$^{-2}$ | 1.63x10$^{-1}$ | 6.10x10$^{-1}$ | 2.09x10$^{-1}$ | 4.13x10$^{-3}$ | $<$ 2x10$^{-16}$ |
| P22 | -2.03x10$^{-2}$ | 1.49x10$^{-1}$ | 8.92x10$^{-1}$ | 1.68x10$^{-1}$ | 4.87x10$^{-3}$ | $<$ 2x10$^{-16}$ |
| P23 | 1.86x10$^{-2}$ | 1.48x10$^{-1}$ | 9.00x10$^{-1}$ | 2.34x10$^{-1}$ | 4.24x10$^{-3}$ | $<$ 2x10$^{-16}$ |
| P24 | 1.67x10$^{-1}$ | 1.66x10$^{-1}$ | 3.14x10$^{-1}$ | 1.67x10$^{-1}$ | 4.89x10$^{-3}$ | $<$ 2x10$^{-16}$ |
| P25 | 3.74x10$^{-1}$ | 1.49x10$^{-1}$ | 1.24x10$^{-2}$ | 2.52x10$^{-1}$ | 4.07x10$^{-3}$ | $<$ 2x10$^{-16}$ |
| P3 | 3.19x10$^{-1}$ | 1.78x10$^{-1}$ | 7.42x10$^{-2}$ | -8.90x10$^{-2}$ | 6.65x10$^{-3}$ | $<$ 2x10$^{-16}$ |
| P4 | 3.14x10$^{-1}$ | 1.90x10$^{-1}$ | 9.73x10$^{-2}$ | -2.14x10$^{-1}$ | 6.64x10$^{-3}$ | $<$ 2x10$^{-16}$ |
| P5 | 3.99x10$^{-1}$ | 1.59x10$^{-1}$ | 1.23x10$^{-2}$ | 6.34x10$^{-2}$ | 4.72x10$^{-3}$ | $<$ 2x10$^{-16}$ |
| P6 | -1.31x10$^{-1}$ | 2.16x10$^{-1}$ | 5.45x10$^{-1}$ | 5.13x10$^{-1}$ | 4.12x10$^{-3}$ | $<$ 2x10$^{-16}$ |
| P8 | 2.57x10$^{-1}$ | 1.40x10$^{-1}$ | 6.73x10$^{-2}$ | 2.83x10$^{-1}$ | 4.30x10$^{-3}$ | $<$ 2x10$^{-16}$ |
| P9 | 2.87x10$^{-1}$ | 1.61x10$^{-1}$ | 7.44x10$^{-2}$ | 2.43x10$^{-1}$ | 4.11x10$^{-3}$ | $<$ 2x10$^{-16}$ |
| T0 | -6.87x10$^{-2}$ | 1.48x10$^{-1}$ | 6.43x10$^{-1}$ | -1.66x10$^{-1}$ | 5.30x10$^{-3}$ | $<$ 2x10$^{-16}$ |
| T1 | 1.19x10$^{-2}$ | 1.56x10$^{-1}$ | 9.39x10$^{-1}$ | -3.33x10$^{-2}$ | 4.64x10$^{-3}$ | 7.40x10$^{-13}$ |
| T10 | -1.79x10$^{-2}$ | 1.41x10$^{-1}$ | 8.99x10$^{-1}$ | 8.55x10$^{-2}$ | 4.10x10$^{-3}$ | $<$ 2x10$^{-16}$ |
| T11 | -8.86x10$^{-2}$ | 1.40x10$^{-1}$ | 5.26x10$^{-1}$ | -1.61x10$^{-1}$ | 4.79x10$^{-3}$ | $<$ 2x10$^{-16}$ |
| T12 | 3.01x10$^{-1}$ | 1.61x10$^{-1}$ | 6.13x10$^{-2}$ | 1.55x10$^{-1}$ | 4.39x10$^{-3}$ | $<$ 2x10$^{-16}$ |
| T13 | 1.44x10$^{-1}$ | 1.49x10$^{-1}$ | 3.33x10$^{-1}$ | 2.03x10$^{-1}$ | 4.15x10$^{-3}$ | $<$ 2x10$^{-16}$ |
| T14 | 1.26x10$^{-1}$ | 1.51x10$^{-1}$ | 4.05x10$^{-1}$ | 7.73x10$^{-2}$ | 4.39x10$^{-3}$ | $<$ 2x10$^{-16}$ |
| T15 | 1.84x10$^{-1}$ | 1.84x10$^{-1}$ | 3.17x10$^{-1}$ | -1.61x10$^{-1}$ | 5.04x10$^{-3}$ | $<$ 2x10$^{-16}$ |
| T16 | 3.29x10$^{-1}$ | 1.46x10$^{-1}$ | 2.45x10$^{-2}$ | 3.79x10$^{-1}$ | 4.94x10$^{-3}$ | $<$ 2x10$^{-16}$ |
| T2 | -1.36x10$^{-1}$ | 1.60x10$^{-1}$ | 3.95x10$^{-1}$ | 2.26x10$^{-1}$ | 4.69x10$^{-3}$ | $<$ 2x10$^{-16}$ |
| T3 | -3.46x10$^{-1}$ | 1.38x10$^{-1}$ | 1.20x10$^{-2}$ | 5.74x10$^{-1}$ | 4.07x10$^{-3}$ | $<$ 2x10$^{-16}$ |
| T4 | -1.28x10$^{-1}$ | 1.50x10$^{-1}$ | 3.95x10$^{-1}$ | -5.11x10$^{-2}$ | 5.89x10$^{-3}$ | $<$ 2x10$^{-16}$ |
| T5 | -4.02x10$^{-1}$ | 1.74x10$^{-1}$ | 2.07x10$^{-2}$ | 6.80x10$^{-1}$ | 5.41x10$^{-3}$ | $<$ 2x10$^{-16}$ |
| T6 | 9.70x10$^{-2}$ | 1.41x10$^{-1}$ | 4.92x10$^{-1}$ | 2.40x10$^{-1}$ | 4.11x10$^{-3}$ | $<$ 2x10$^{-16}$ |
| T7 | -2.07x10$^{-1}$ | 1.35x10$^{-1}$ | 1.24x10$^{-1}$ | 1.59x10$^{-1}$ | 4.42x10$^{-3}$ | $<$ 2x10$^{-16}$ |
| T8 | 8.55x10$^{-2}$ | 1.50x10$^{-1}$ | 5.68x10$^{-1}$ | 5.83x10$^{-1}$ | 4.46x10$^{-3}$ | $<$ 2x10$^{-16}$ |
| T9 | 3.88x10$^{-2}$ | 1.50x10$^{-1}$ | 7.96x10$^{-1}$ | -1.28x10$^{-2}$ | 4.46x10$^{-3}$ | 4.05x10$^{-3}$ |
| R-squared | 1.00x10$^{-1}$ | | | 4.10x10$^{-1}$ | | |
| Log-likelihood | 1.03x10$^{+4}$ | | | | | |
| phi | 5.30x10$^{-1}$ | 8.31x10$^{-3}$ | $<$ 2x10$^{-16}$ | | | |
| F-statistic | | | | 4.59x10$^{+4}$ | | |
| F-statistic DF | | | | 6.80x10$^{+1}$ | | |
